# Differential effects of MATR3 variants on its cryptic splicing repression function

**DOI:** 10.1101/2023.12.07.570685

**Authors:** Mashiat Khan, Xiao Xiao Lily Chen, Michelle Dias, Jhune Rizsan Santos, Sukhleen Kour, Justin You, Rebekah van Bruggen, Mohieldin M.M. Youssef, Ying-Wooi Wan, Zhandong Liu, Jill A. Rosenfeld, Qiumin Tan, Udai Bhan Pandey, Hari Krishna Yalamanchili, Jeehye Park

**Affiliations:** Department of Molecular Genetics, University of Toronto, Toronto, M5S 1A1, Canada; Peter Gilgan Centre for Research and Learning, The Hospital for Sick Children, Toronto, M5G 0A4, Canada; Department of Pediatrics, Baylor College of Medicine, Houston, 77030, USA; Jan and Dan Duncan Neurological Research Institute at Texas Children’s Hospital, Houston, 77030, USA; Department of Pediatrics, Children’s Hospital of Pittsburgh, University of Pittsburgh Medical Center, Pittsburgh, 15224, USA; Department of Molecular and Human Genetics, Baylor College of Medicine, Houston, 77030, USA; Baylor Genetics Laboratories, Houston, 77021, USA; Department of Cell Biology, University of Alberta, Edmonton, T6J 2H7, Canada; Department of Human Genetics, University of Pittsburgh, School of Public Health, Pittsburgh, 15260, USA; USDA/ARS Children’s Nutrition Research Center, Department of Pediatrics, Baylor College of Medicine, Houston, Texas, 77030, USA

**Keywords:** Cryptic splicing, MATR3, RNA-binding protein, amyotrophic lateral sclerosis, neurodevelopmental disease

## Abstract

MATR3 is an RNA-binding protein implicated in neurodegenerative and neurodevelopmental diseases. However, little is known regarding the role of MATR3 in cryptic splicing within the context of functional genes and how disease-associated variants impact this function. We show that loss of MATR3 leads to cryptic exon inclusion in many transcripts. We reveal that ALS-linked S85C pathogenic variant reduces MATR3 solubility but does not impair RNA binding. In parallel, we report a novel neurodevelopmental disease-associated M548T variant, located in the RRM2 domain, which reduces protein solubility and impairs RNA binding and cryptic splicing repression functions of MATR3. Altogether, our research identifies cryptic events within functional genes and demonstrates how disease-associated variants impact MATR3 cryptic splicing repression function.

## Introduction

Alternative splicing plays a critical role in enhancing the diversity and complexity of the transcriptome and proteome generated from a limited number of genes in eukaryotes [1]. This process is particularly prolific in the nervous system, enabling a greater degree of functional complexity [2]. The regulation of splicing is orchestrated through the interplay of cis-acting elements—like the splicing consensus sequences found in pre-mRNA—and trans-acting elements, including the foundational spliceosome machinery. In addition, RNA-binding proteins act as regulatory players that enhance or suppress splicing. RNA-binding proteins have RNA binding domains, which recognize specific RNA binding motifs. There are more than 1,000 RNA-binding proteins in the human genome with the vast majority expressed in the nervous system [3]. However, how these RNA-binding proteins regulate splicing under physiological and pathogenic conditions remains largely unexplored.

While alternative splicing is the canonical splicing of previously annotated exons [4], cryptic splicing is the non-canonical splicing of previously unannotated exons or inclusion of novel non-conserved intronic sequences, a process that generally occurs with low efficiency and is often considered erroneous [5]. Several RNA-binding proteins have recently been identified to play a role as a repressor of cryptic splicing [6–9]. Loss of a cryptic splicing repressor can result in enhanced inclusion of the cryptic exon, often leading to frameshift mutations and/or premature termination and subsequent destabilization of the transcript or production of an aberrant protein [5,10,11]. These aberrant cryptic splicing events have been observed in neurodegenerative diseases, including Alzheimer’s disease (AD), frontotemporal dementia (FTD) and amyotrophic lateral sclerosis (ALS) [9–12]. TAR DNA-binding protein 43 (TDP-43) is an FTD/ALS-linked RNA-binding protein that plays a major role as a cryptic splicing repressor [9,13,14]. Recent studies demonstrated that loss of TDP-43 exposes the cryptic splicing sites in an ALS risk factor gene *UNC13A* [14,15], providing additional support for the role of TDP-43 as a cryptic splicing repressor. In addition to TDP-43, there are a handful of RNA-binding proteins that are associated with ALS [16], but little is known about whether other ALS-linked RNA-binding proteins including Matrin 3 (MATR3) are involved in cryptic splicing repression within functional genes.

MATR3 is a DNA and RNA-binding protein originally found as a component of the nuclear matrix in rat liver nuclei [17]. MATR3 is predominantly localized to the nucleus, found diffusely in the nucleoplasm and enriched on the nuclear scaffold [18,19]. MATR3 contains an N-terminal nuclear export signal (NES), a C-terminal bipartite nuclear localization signal (NLS), two RNA recognition motif (RRM) domains and two CCHH-type zinc finger (ZF) domains [18,20,21]. The ZF domains have been shown to interact with DNA, while the RRM domains bind RNA [18]. MATR3 has been implicated in DNA-regulating activities, including chromosomal organization and transcription, as well as RNA-regulating activities, including mRNA export, mRNA stability, RNA splicing and processing, and the nuclear retention of hyper-edited mRNA [18,19,22–28].

As a regulator of RNA splicing, MATR3 binds pyrimidine-rich intronic sequences [24,25]. Most of the alternative splicing events regulated by MATR3 are exon skipping events [24,25], indicating that the major splicing function of MATR3 is to repress exon inclusion. In addition, MATR3 interacts with polypyrimidine tract binding protein 1 (PTBP1), another RNA-binding protein that acts as a splicing repressor by binding pyrimidine-rich motifs in intronic regions [24,29]. While many splicing events are shared with PTBP1, the majority of the events are regulated by MATR3 alone in a PTBP1-independent manner [24,25]. In addition, MATR3 works alongside PTBP1 to prevent the inclusion of cryptic exons within long interspersed nuclear elements (LINEs) [30]; however global MATR3-dependent cryptic splicing events within the context of functional genes have not been reported. Furthermore, the molecular mechanism by which MATR3 regulates cryptic splicing is still unclear.

Missense variants in *MATR3* have been identified in familial ALS cases as well as in sporadic cases [31–36]. These ALS-linked variants are located outside of the functional domains, largely clustered in the N-terminus or the C-terminus of the intrinsically disordered regions of MATR3 [16]. The S85C pathogenic variant is the most common; first linked to familial distal myopathy, but later reclassified as familial ALS [31,37,38]. Accumulating studies suggest that the S85C pathogenic variant leads to the loss of MATR3 function [39–41]. However, how the S85C pathogenic variant alters MATR3 properties and function and contributes to ALS pathogenesis remains unclear. Intriguingly, a recent study reported that a missense variant in *MATR3* is implicated in neurodevelopmental disorders [42]. A child presenting with severe neurodegeneration, muscular hypotonia and epileptic seizures was found to carry a *de novo* E436K variant, which resides in the RRM1 domain of MATR3 [42]. How these neurodevelopmental disease-linked variants impair the molecular functions of MATR3 and how these mechanisms differ from those caused by ALS-linked variants, and thereby lead to varying clinical outcomes, remains undetermined.

In this study, we investigate the role of MATR3 in cryptic splicing within functional genes and investigate the mechanism by which this occurs. We identify global cryptic splicing events within functional genes upon MATR3 knockdown and show repression of these events upon MATR3 overexpression. We demonstrate that the RRM2 domain of MATR3 is required for its cryptic splicing function. We also show how disease-associated variants impair the cryptic splicing and RNA binding functions of MATR3. Our results show that the N-terminal S85C pathogenic variant slightly impairs the cryptic splicing function of MATR3 and decreases the solubility of MATR3 but does not impair its RNA binding ability. We identified a novel heterozygous M548T variant in MATR3 in an individual with neurodevelopmental phenotypes and mitochondrial complex III deficiency. We demonstrate that the M548T variant, located in the RRM2 domain, leads to a loss of solubility, cryptic splicing repression ability, and RNA binding ability. Together, our data demonstrate that MATR3 is involved in repressing the inclusion of cryptic exons and that the RRM2 domain is required for this function. Furthermore, our findings reveal that the two disease-associated variants differentially affect MATR3 properties and/or splicing repression ability, which may help explain the two different consequences, neurodegeneration and neurodevelopmental defects.

## Materials and Methods

### RNA sequencing

HeLa cells were seeded in 6-well plates the day before transfection at a confluency of around 30-40% while SH-SY5Y cells were seeded at a confluency of 50%. The next day, the cells were transfected with 2 µg of scrambled siRNA (MISSION siRNA Universal Negative Control #2 Sigma-Aldrich SIC002-5x1nmol) or 2 µg of MATR3 siRNA (MISSION esiRNA human MATR3 Sigma-Aldrich EHU113801-50UG) using Lipofectamine 3000 Transfection Reagent (ThermoFisher Scientific L3000015) as per manufacturer’s instructions. MATR3 siRNA was used to knock down MATR3 with scrambled siRNA as a negative control. The cells were harvested 72 hours after transfection and RNA was extracted. The Centre for Applied Genomic (TCAG) prepared RNA libraries from the total rRNA depletion RNA. The RNA was pair-end sequenced. An average of 90 million read pairs per sample were sequenced, with a read length of 150bp.

### RNA sequencing data analysis

Sequencing quality and adapter contamination were assessed using FastQC v0.10.1 [43]. Adapter sequences were trimmed using fastp v0.20.0 [44]. Trimmed reads were aligned to human reference genome GRCh38 using HISAT2 [45]. Raw FASTA sequence and annotations of genome build GRCh38 release 24 were downloaded from GENCODE portal. Raw genome was indexed using HISAT2-build command. Alignments were saved in Sequence Alignment/Map (SAM) format. Coordinate sorted binary format (BAM) files were generated using SAMtools [46]. The RNA seq results are deposited in the NCBI GEO (GSE number GSE205343).

### Differentially Expressed Genes (DEGs)

Gene expression values from each sample were quantified as the number of reads mapped (to a specific gene) using FeatureCounts [47]. Genes with an average read count < 50 across the samples were considered not expressed and were excluded from the DEG analysis. Raw read counts were normalized and then tested for differential expression using *DESeq2* [48]. A false discovery rate (FDR) cut-off of 0.05 and a fold change cut-off of 20% (-0.263 ≤ log_2_(FC) ≥+0.263) were used to call DEGs.

### Alternative Splicing analysis

Alternative splicing events were quantified and classified using *rMATS* [49]. Alignment files (BAM) and reference annotations (GTF) from NCBI Reference Sequence database were used. rMATS classified splicing events into 5 categories: skipped exons, retained introns, mutually exclusive exons, alternative 5ʹ and 3ʹ splice sites. An FDR cut-off of 0.05 and an inclusion level difference cut-off of less than -0.2 or greater than 0.2 were used to screen for statistically significant changes.

### Cryptic Splicing analysis

We used CrypSplice [8,50] to infer novel cryptic splice sites. CrypSplice evade full-length isoform and ASM quantifications and is based only on junction counts. RNA-Seq data is best abstracted as the number of reads mapped to a specific feature, here we map exon-exon junctions. Junctions were quantified as the ratio of junction reads to the 5ʹ splice site coverage in respective conditions and subjected to a beta binomial test. Reference annotations (GTF) from NCBI Reference Sequence database were used. Junctions with an FDR of less than 0.05 were called significant. We updated CrypSplice to V1.7 and the additional POverA-M parameters were set to 1, 10 and 10 respectively.

### Sequence composition analysis

FASTA sequences with a window of 50 nucleotides into the intron and 25 nucleotides into the exon were extracted from 3ʹ end of the cryptic junction with FDR ≤ 0.05 and junction difference > 0 using bedtools [51]. Sequences from the reverse were reverse complimented. Nucleotide frequencies and sequence logo were computed using WebLogo [52].

### CLIP-Seq analysis

MATR3 CLIP-Seq data was downloaded from ARRAYEXPRESS (E-MTAB-3107). Raw read quality was assessed using FastQC [43]. Reads were trimmed using Trimmomatic [53] and aligned to human reference genome GRCh38 using STAR. Alignment files were normalized and converted to bigwig format using deeptools [54]. For the illustration, the sample with the best sequencing quality and mapping (Matr3_iCLIP_medRNase_repl1) was used.

### Premature termination codon (PTC) analysis

We employed the detection of premature termination codons (PTCs) as an indicating marker to evaluate the propensity of cryptic splice junctions to induce nonsense-mediated decay (NMD). We first constructed all possible transcript structures from each of the MATR3 KD samples using StringTie [55]. Subsequently, we extracted the transcript sequences with gffread [56]. We then searched for potential in-frame stop codons and calculated the distance from the second last in-frame stop codon (with the last one being the actual stop codon) to the transcript’s last exon-exon junction. Only transcripts containing at least two in-frame stop codons were included in this analysis. Transcripts with the PTC positioned more than 55 nt away from the last exon-exon junction are identified as NMD-prone. Lastly, to infer the NMD potential of cryptic junctions, we aligned them with the identified NMD-susceptible transcripts from individual MATR3 KD samples.

### Cell lines

Immortalized HeLa cells and SH-SY5Y cells were cultured in Dulbecco’s modified Eagle’s medium (DMEM, Wisent 319-005-CL) with 10% fetal bovine serum (Wisent 080-150) and maintained at 37°C in a humidified incubator containing 5% CO_2_.

*MATR3 ^S85C/+^*iPSCs were generated by iPSC Neurodegenerative Disease Initiative (iNDI) and were a generous gift from Dr. Michael E. Ward (NIH/NINDS) [57,58]. The control isogenic Kolf2.1 cultured was a kind gift from Professor William C. Skarnes (The Jackson Laboratory). The iPSCs were maintained and propagated in mTESR1 (STEMCELL technologies) on Matrigel (Corning) coated plates.

### iPSCs cells culture and differentiation into neurons

*MATR3 ^S85C/+^*and isogenic Kolf2.1 control iPSCs were differentiated into neurons following the previously described protocol [59–61]. Briefly, iPSCs were dissociated with ReLeSR™ (STEMCELL technologies) and grown to 80–90% confluency on Matrigel coated plates in mTESR1 medium. The iPSCs were then cultured for 7 days in Neurobasal/DMEM-F12 medium (1:1 v/v) containing 2% B27 (Gibco, 17054–044), 1% N2 (Gibco, 17502–048), 1% Glutamax (Gibco), and 1% non-essential amino acids (NEAA, Gibco, 11140050) along with molecules essential for generation of neuro-progenitor cells such as smoothened agonist (SAG, Cayman chemicals 11914), SB431542 (STEMCELL technologies), LDN (Sigma SML0559), and retinoic acid (RA, Sigma R2625). For the next seven days, the cells were switched to N2-B27-neurobasal-media supplemented with DAPT (Cayman, 13197), SU5406 (Cayman, 131825), RA and SAG. The neuro-progenitor cells were then dissociated with DNase I (Worthington) supplemented TrypLE Express (GIBCO) and cultured on poly-ornithine and laminin-coated plates in N2-B27-neurobasal media containing growth factors: BDNF, GDNF, CNTF (Peprotech) and ascorbic acid (Sigma, A4403). The differentiated neurons were grown for 7 days and processed for subsequent RNA and protein extraction.

### Plasmid constructs

The vector backbone 1436 pcDNA3 Flag HA was a gift from Dr. William Sellers (Addgene plasmid # 10792). MATR3 WT and the single domain deletion constructs were gifts from Dr. Yossi Shiloh (Addgene plasmids #32880, 32881, 32882, 32883, 32884). We generated the double RRM domain deletion mutant and the double ZF domain deletion mutant using Gibson assembly, and then inserted into the pcDNA3 backbone via restriction enzyme cloning using BamH1 (New England Biolabs R3136L) and Xho1 (New England Biolabs R0146L). MATR3 M548T was generated via site-directed mutagenesis with QuikChange II (Agilent Technologies 200524, forward primer: TTCAAACAATGGTCAACCGTTGCCATTGCATCTTCTCTTG, reverse primer: CAAGAGAAGATGCAATGGCAACGGTTGACCATTGTTTGAA). MATR3 S85C was generated via site-directed mutagenesis with QuikChange II (Agilent Technologies 200524, forward primer: TACTTCTTCCCATAATTTGCAGTGTATATTTAACATTGGAAGTAGAG, reverse primer: CTCTACTTCCAATGTTAAATATACACTGCAAATTATGGGAAGAAGTA).

### Transfection

For rescue experiments, HeLa cells were seeded the day before transfection into 6-well plates. The next day, they were transfected at 40% confluency with 200 ng of the pcDNA-MATR3 vectors and 10 μM of either control or MATR3 siRNA per well using the Lipofectamine 3000 transfection reagent (ThermoFisher Scientific L3000015). ON-TARGETplus non-targeting siRNA #1 ((Dharmacon D- 001810-01-05), ON-TARGETplus non-targeting siRNA #2 (Dharmacon D-001810-02-05) and custom MATR3 siRNA targeting the 3ʹ UTR (Dharmacon, sense sequence: GCAAGGAGAUUUAAUGAUUUU, antisense sequence: AAUCAUUAAAUCUCCUUGCUU) were used. The media was changed 24 hours post-transfection.

For RNA immunoprecipitation experiments, HeLa cells were seeded the day before transfection into 6-well plates. The next day, they were transfected at 70% confluency with 1 ug of the pcDNA-MATR3 per well, two wells per construct, using the Lipofectamine 3000 transfection reagent. The media was changed 24 hours post-transfection.

For soluble/insoluble fractionation experiments, HeLa cells were seeded the day before transfection into 6-well plates. The next day, they were transfected at 70% confluency with 500 ng of the pcDNA-MATR3 vectors per well using the Lipofectamine 3000 transfection reagent. The media was changed 24 hours post-transfection.

### RNA extraction

Cells were harvested for RNA isolation 48-hours post-transfection using TRIzol reagent (ThermoFisher Scientific 15596018) following manufacturer’s instructions. Briefly, each well was washed with D-PBS (Wisent 311-425-CL), then resuspended in TRIzol reagent (ThermoFisher Scientific 15596018) prior to transferring to a centrifuge tube. Chloroform was added and the samples were centrifuged to separate the RNA-containing aqueous phase from the protein-containing organic phase. RNA was precipitated using isopropanol, washed in 75% ethanol, dried, resuspended in DEPC-treated water (ThermoFisher Scientific 4387937), and kept at -80°C for long-term storage.

For RNA immunoprecipitation experiments, RNA was extracted from the RNA input samples (thawed on ice) and RNA IP samples as described above, except for the addition of RNA grade glycogen (ThermoFisher Scientific R0551) prior to RNA precipitation, used to increase the RNA pellet visibility. RNA was resuspended in DEPC-treated water (ThermoFisher Scientific 4387937) and stored on ice for cDNA generation.

### cDNA synthesis

For rescue experiments, cDNA was generated from previously extracted RNA. RNA concentrations were measured using the Thermo Scientific Nanodrop spectrophotometer. Using a starting concentration of 2 μg, cDNA was generated from RNA using random hexamers (ThermoFisher Scientific N80801270 and M-MLV reverse transcriptase kit (Invitrogen 28025-013) with the addition of RNaseOUT recombinant ribonuclease inhibitor (ThermoFisher Scientific 10777-019) as per manufacturer’s instructions. cDNA was kept at -20 °C for long-term storage.

For RNA immunoprecipitation experiments, RNA input and IP sample concentrations were measured using the Thermo Scientific Nanodrop spectrophotometer. The maximum amount of cDNA possible was generated based on the lowest measured concentration.

### RT-PCR

For rescue experiments, RT-PCR was performed on previously generated cDNA (8 ng) using KOD Hot Start DNA Polymerase (VWR CA80511-386). The primer sequences for the target transcripts and number of cycles are outlined in **Table 3**. Cycle numbers were kept at 32 cycles and below to ensure reactions were in the exponential amplification state, and saturation controls were used while imaging to ensure the accuracy of subsequent quantification. *GAPDH* was run as a loading control. The expression of the MATR3 vectors was evaluated using the *MATR3* internal primers (F4/R4 was used for the ZF mutants, F5/R5 was used for all other vectors). Knockdown of endogenous *MATR3* was evaluated with 5ʹUTR primers as the UTRs are absent from the vectors. For *UQCRC2*, *PHF20, CACNB2*, and *UHRF2*, inclusion of the cryptic exon was evaluated with the cryptic exon primers, and the total levels of the targets was evaluated with the internal primers.

For RNA immunoprecipitation experiments, RT-PCR for *UQCRC2* and *CACNB2* total transcript levels was performed on previously generated RNA input and RNA IP cDNA using KOD Hot Start DNA Polymerase (VWR CA80511-386). For *UQCRC2*, approximately 10 ng of cDNA was used in each reaction along with the *UQCRC2* internal primers (**Table 3**). For *CACNB2*, approximately 10 ng of cDNA was used for the input reactions and approximately 40ng of cDNA was used for the IP samples, along with the *CACNB2* internal primers (**Table 3**).

### Quantitative PCR (qPCR)

qPCR was run with cDNA previously generated from rescue experiments in a 10 µl reaction using iTaq Universal SYBR Green Master Mix (BioRad 1725121) with readout using the Applied Biosystems ViiA 7 real-time PCR system with standard cycling parameters (50°C for 1 min, 95 °C for 1.25 min, 40 cycles of 95 °C for 15s and 60 °C for 60 s), followed by standard melt curve analysis (95 °C for 15 s, 65 °C to 95 °C at 0.3 °C/s). ΔΔCt was calculated with the housekeeping gene *GAPDH* as reference and siCTRL GFP or siCTRL empty vector samples as the calibrator. The relative expression level of the targets was calculated using the 2^-ΔΔCt^ method. The internal and cryptic exon primers used in the qPCR reaction are outlined in **Table 3**. Statistical analyses were performed on Graphpad Prism Version 9.3.0, and significance was determined using Welch’s t-test.

### RNA immunoprecipitation (RIP)

The day before starting the IP, Protein G Mag Sepharose beads (Sigma-Aldrich GE28-9513-79) were washed in cell lysis buffer and blocked in 10% BSA (Jackson ImmunoResearch 001-000-162) at 4°C. Beads were divided into two aliquots per sample, with one sample stored on ice (referred to hereafter as pre-cleared beads) and the other incubated with mouse anti-FLAG (referred to hereafter as IP beads) for 3-4 hours at 4°C. Cells were harvested for IP 48-hours post-transfection. Each well was washed with D-PBS (Wisent 311-425-CL), trypsinized and then resuspended in DMEM supplemented with 10% FBS, pooling wells transfected with the same construct. Samples were divided into two for total RNA analysis and immunoprecipitation. The RNA samples were resuspended in TRIzol reagent and stored at -80°C overnight to be used as the RNA input samples. The IP samples were resuspended in cell lysis buffer, incubated on ice for 30 minutes, then centrifuged at 13,250 rpm for 20 minutes at 4°C. For input samples, 50 μL of the soluble supernatant was transferred to a different tube, 4X BOLT LDS loading dye and 10X BOLT reducing buffer were added to each sample to a concentration of 1X, which were then boiled at 80°C for 10 minutes, then placed at -20°C for storage. The remaining soluble supernatant was transferred from the IP samples to a new tube containing 10 μL of pre-clear beads and incubated for 1 hour at 4°C on a rotator for pre-clearing. After pre-clearing, the protein concentration was measured using a Bradford assay (Sigma-Aldrich/Millipore B6816-500ML) as per manufacturer’s instructions in optically clear 96-well plate (VWR 62402-933) on the Molecular Devices VersaMax 190 plate reader. BSA protein standards (0-2mg/mL) were used as a control. To normalize protein concentrations, all samples were diluted to the lowest measured protein concentration. These normalized samples were then added to the IP beads, and incubated overnight on a rotator at 4°C. The next day, the supernatant from the samples was discarded, and the IP beads were washed and then divided into two aliquots. One aliquot was resuspended in TRIzol reagent. The other aliquot was resuspended in 4X BOLT LDS loading dye and 10X BOLT reducing buffer, boiled at 80°C for 10 minutes, then placed at -20°C to be used as the IP samples for Western blot analysis.

### Protein extraction

For HeLa cells, cells were harvested for protein analysis 48-hours post-transfection. Each well was washed with D-PBS (Wisent 311-425-CL), then resuspended in cell lysis buffer (50 mM Tris pH 7.4, 150 mM NaCl and 0.5% Triton-X 100, supplemented with PhosStop phosphatase inhibitor (Roche 04906845001) and cOmplete protease inhibitor (Sigma-Aldrich/Millipore 5056489001), referred to hereafter as cell lysis buffer). Samples were incubated on ice for 30 minutes, then centrifuged at 13,250 rpm for 20 minutes at 4°C. The soluble supernatant (soluble fraction) was transferred to a new tube. The pellet was resuspended in urea buffer (7M Urea, 2M Thiourea, 4% CHAPS, 30mM TrisHCl pH8.5, supplemented with PhosSTOP phosphatase inhibitor and cOmplete protease inhibitor), sonicated at 20Amps for 5 pulses (1 sec on, 1 sec off) using the Misonix Sonicator S-4000 (insoluble fraction)., Samples were then mixed with 4X BOLT LDS sample buffer (ThermoFisher Scientific B00008) and 10X BOLT reducing buffer (ThermoFisher Scientific B0004). Soluble fractions were boiled at 80°C for 10 minutes while insoluble reactions were incubated at room temperature for 10 minutes, then placed at -20°C for storage.

### Western blots

Samples were loaded onto an 8% 1.5 mm acrylamide gel. For RNA immunoprecipitation experiments, 5% of the input and 50% of the IP were loaded. Gels were transferred onto a nitrocellulose membrane at 320 mA for 2 hours at 4°C. Membranes were cut between the 50 and 64 kDa markers and incubated in Li-Cor Intercept blocking buffer (Mandel Scientific Company LIC-927-60003) diluted 1:1 in 1X TBS for 1 hour at room temperature, then incubated in primary antibody overnight at 4°C. The lower half of the membrane was probed with mouse anti-GAPDH (Calbiochem clone 6C5 CB1001-500UG) at 1:10,000 while the upper half was probed with mouse anti-FLAG (Sigma-Aldrich/Millipore F1804-1MG) at 1:1000. The next day, membranes were washed in 1X TBS at room temperature, then incubated in secondary antibody (Li-Cor IRDye 800CW Goat anti-Mouse IgG, Mandel Scientific 925-32210, 1:10,000) for 1 hour at room temperature. Membranes were then washed 1X TBS at room temperature and imaged on the Li-Cor Odyssey Fc imager.

### Quantification and statistical analyses

For rescue experiments, agarose gel images were quantified using ImageJ 1.53f5. The relative density of the cryptic exon of each target was normalized to the relative density of *GAPDH* for each corresponding sample. Statistical analyses were performed on Graphpad Prism Version 9.3.0. Normalized relative density measurements were imported and significance was determined by Welch’s t-test.

For RNA immunoprecipitation experiments, agarose gel and western blot images were quantified using ImageJ 1.53f5. The relative density of each target transcript on the agarose gel was normalized to the relative density of the corresponding sample on the blot to determine the binding ratio of the target transcript to the protein. Statistical analyses were performed on Graphpad Prism Version 9.3.0. Ratio of binding measurements were imported, and significance was determined by Welch’s t-test.

Western blot images were quantified using ImageJ 1.53f5. The relative density of MATR3 was normalized to the relative density of GAPDH in the corresponding sample. Statistical analyses were performed on Graphpad Prism Version 9.3.0. Normalized relative density measurements were imported and significance was determined by Welch’s t-test.

### Cycloheximide treatment in HeLa cells

HeLa cells were seeded the day before transfection into 6-well plates. The next day, they were transfected at 40% confluency with 10 μM of either control or MATR3 siRNA per well (see Transfection section for details). The media was changed 24 hours post-transfection. After 48 hours of transfection, cells were treated with either DMSO (to a final concentration of 1%) or cycloheximide (to a final concentration of 30 µg/mL) for 4.5 hours. Cells were then harvested for RNA and RT-PCRs were performed from generated cDNA (see RNA extraction, cDNA generation and RT-PCR sections for details).

### Cryptic splicing analysis in iPSC neurons

Total RNA was isolated from differentiated *MATR3^S85C/+^* and control isogenic Kolf2.1 neurons by using the PureLinkTM RNA mini kit (Invitrogen), by following the manufacturer’s instructions. cDNA was synthesized from 500 ng of total RNA by using PCR supermix (Invitrogen^TM^) and 1 mM MgSO_4_. The PCR reaction conditions, and number of cycles were performed similarly to RT-PCR in HeLa cells. The internal and cryptic exon primers used in the PCR reaction are outlined in **Table 3**. The amplified DNA was separated by using 6% TBE gels (Invitrogen) and imaged.

### Protein analysis in iPSC neurons

For separating the soluble protein fractionation, *MATR3^S85C/+^* and control isogenic Kolf2.1 neurons were lysed in NP40 lysis buffer (0.5% NP40, 10 mM Tris HCl pH 7.8, 10 mM EDTA, 150 mM NaCl, 2.5 mM Na orthovanadate) containing 1x protease inhibitor (Invitrogen^TM^). The lysate was centrifuged at 21,000 x g for 30 minutes and the supernatant (soluble fraction) was collected and boiled in 1X NuPage™ LDS-Sample buffer (Invitrogen, NP0007) for 2 minutes. The soluble fraction was separated using 3-8% Tris-acetate NuPage gel (Novex/Life Technologies) and the proteins were transferred onto a nitrocellulose membrane (Invitrogen IB23001). Following incubation in 2.5% QuickBlocker reagent (EMB Millipore WB57-175GM) for an hour at room temperature, the membrane was probed with mouse anti-MATR3 (Abcam, 1:2000) and mouse anti-tubulin (Sigma-Aldrich, 1:10,000) antibodies at 4 °C overnight. The next day, the membrane was incubated with goat anti-mouse 680 (Invitrogen, 1:10,000) secondary antibody for an hour at room temperature followed by washes with 1% Tween-20 in Tris buffered saline (TBST). The membrane was imaged using Odyssey® CLx (LI-COR Biosciences) and the integrated band densities (IDVs) were calculated using LI-COR image studio software.

## Results

To identify the gene expression changes that occur upon loss of MATR3, we performed RNA sequencing on HeLa cells treated with either MATR3 siRNA or control siRNA (confirmation of knockdown on Supplementary Figure S1), then performed differential gene expression analysis using DESeq2. After filtering for an adjusted P-value ≤ 0.05 and log_2_ fold change < -0.263 or > 0.263, we found a total of 357 differentially expressed genes with 176 upregulated genes and 181 downregulated genes upon MATR3 knockdown (Supplementary Figure S2, Supplementary Data File 1). We then proceeded to investigate the splicing changes that occur upon MATR3 knockdown. To determine the alternative splicing events affected by MATR3 depletion, we used replicative multivariate analysis of transcript splicing (rMATS) to look for differential splicing events. After filtering with a false discovery rate (FDR) ≤ 0.05 and an inclusion level difference greater than 0.2 or less than -0.2, we identified a total of 2,131 alternative splicing events upon MATR3 knockdown compared to the control (Supplementary Figure S3, Supplementary Data File 2). The largest category of alternative splicing events affected by MATR3 knockdown was cassette exons (Supplementary Figure S3). We saw increased inclusion of these events upon MATR3 knockdown, suggesting that the major role of MATR3 is a splicing repressor, which is in accordance with previous reports [24,25].

To examine whether MATR3 plays a role in cryptic splicing within functional genes, we used CrypSplice [8] to analyze global cryptic splicing events upon MATR3 depletion. We focused on splicing events with an adjusted P-value ≤ 0.05 and found a total of 899 cryptic splicing events upon MATR3 knockdown (Figure 1A, Supplementary Data File 3), with the greatest number of events falling into the “Novel Acceptor” category (Figure 1A). Functional analysis of the 899 significant cryptic splicing events indicated that the cryptic splicing changes are enriched in genes that encode proteins involved in intracellular transport (Supplementary Figure S4A). It was previously reported that MATR3 binds to pyrimidine-rich intronic regions [25]. Sequence composition analysis revealed the presence of pyrimidine-rich intronic regions towards the 3’ end of the cryptic junction gains (Figure 1B), suggesting the presence of MATR3 binding sites adjacent to cryptic events. This polypyrimidine tract upstream of the exon is also a consensus 3’ splice site element in general and thus is not enough to decipher the contributory role of MATR3. Therefore, we analyzed CLIP-Seq data from HeLa cells generated by a previous study [24] and found MATR3 binding within or near these cryptic exons (Figure 1C-D, Supplementary Figure S4B-S4C, Supplementary Figure S10A, shown in yellow). Intronic pyrimidine-rich regions (Figure 1B) and MATR3 binding (Figure 1C-D, Supplementary Figure S4B-S4C, Supplementary Figure S10A) within or near the cryptic exons strongly suggest the role of MATR3 in cryptic splicing. Focusing on splicing events with > 10% cryptic exon inclusion (junction strength difference) upon MATR3 knockdown, we identified 143 distinct cryptic splicing events across 122 genes. To validate these events, we chose five candidate genes containing cryptic splicing events with high significance and inclusion levels or of particular relevance to ALS (Table 1): *UQCRC2*, which encodes a subunit of mitochondrial electron transport chain complex III (Figure 1C); *PHF20,* which encodes a component of the MOF (male absent on the first) histone acetyltransferase complex (Supplementary Figure S4B); *CACNB2*, which encodes the calcium voltage-gated channel auxiliary subunit beta 2 (Supplementary Figure S4C); *UHRF2,* which encodes an E3 ubiquitin ligase involved in the maintenance of DNA methylation and has been previously shown to play a role in the intranuclear degradation of polyglutamine aggregates (Figure 1D) [62–67]; and *ADARB1*, which encodes an adenosine deaminase that catalyzes the conversion of adenosine to inosine in double stranded RNAs and has been previously shown to display pathology in ALS patients (Supplementary Figure S10A) [68,69]. Next, we validated the inclusion of these cryptic exons upon MATR3 knockdown using RT-PCR in both HeLa cells and SH-SY5Y cells, a human neuronal-like cell line (Supplementary Figure S5-S7). These results corroborate that these cryptic splicing events are regulated by MATR3.

**Figure 1.**
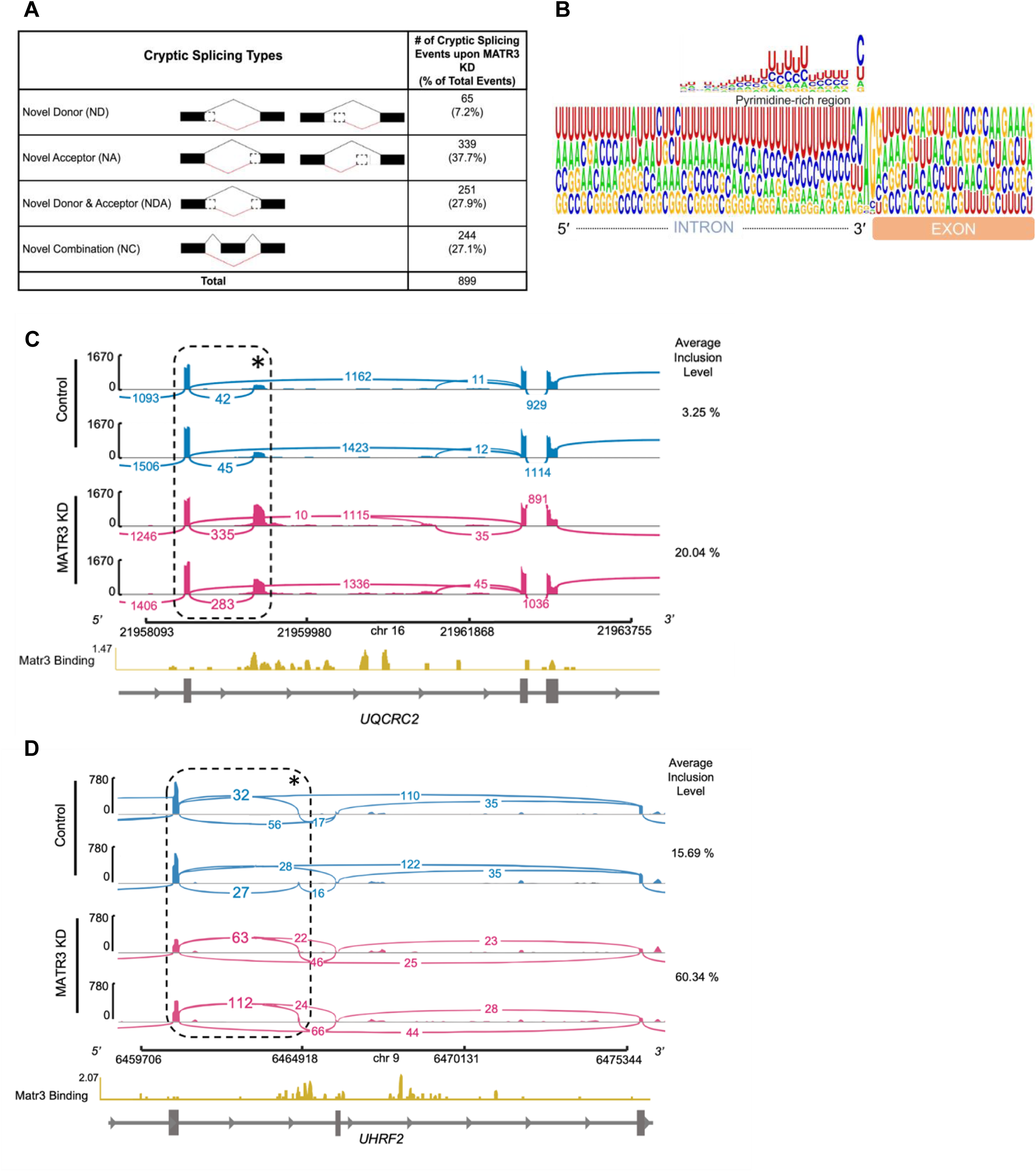
Knockdown of MATR3 results in cryptic splicing events. **A.** Tabulated result of the significant cryptic splicing events found in MATR3 KD summarized by category: novel donor (ND), novel acceptor (NA), novel donor and acceptor (NDA) or novel combination of known acceptors and donors (NC). Significant events were identified using an adjusted P-value ≤ 0.05. **B.** Sequence composition showing pyrimidine-rich sequences in the intronic region towards the 3’ end of the cryptic junctions with an FDR ≤ 0.05 and junction difference > 0. **C-D.** Sashimi plot generated using IGV genome browser illustrating the read coverage for two biological replicates of control (blue) and MATR3 knockdown (pink) for a region of *UQCRC2* and *UHRF2* transcribed from left to right. The bar graph denotes read depth, and the arcs represent the number of reads connecting each splice junction, with the junction coverage minimum set to 10. The region within black dashed lines represents the cryptic junction of interest with the cryptic exon marked by an asterisk, and inclusion levels of the cryptic exon within this region are shown as percentages to the right of the tracks. MATR3 binding analyzed from CLIP-Seq data from Coelho *et al.,* 2015 is shown in yellow.

**Table 1.**
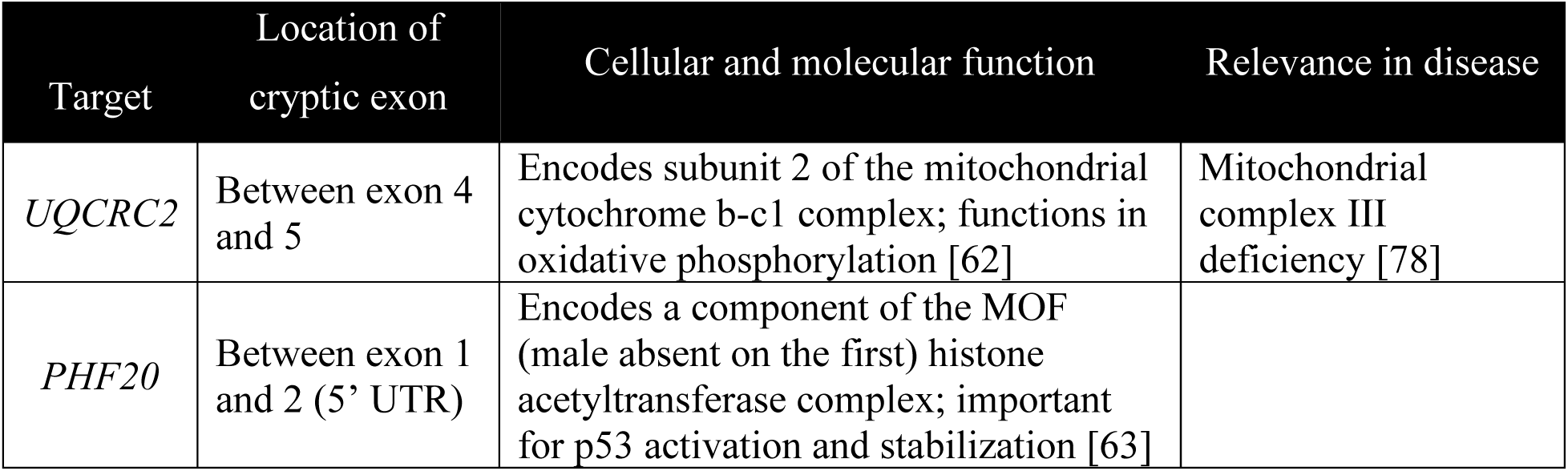
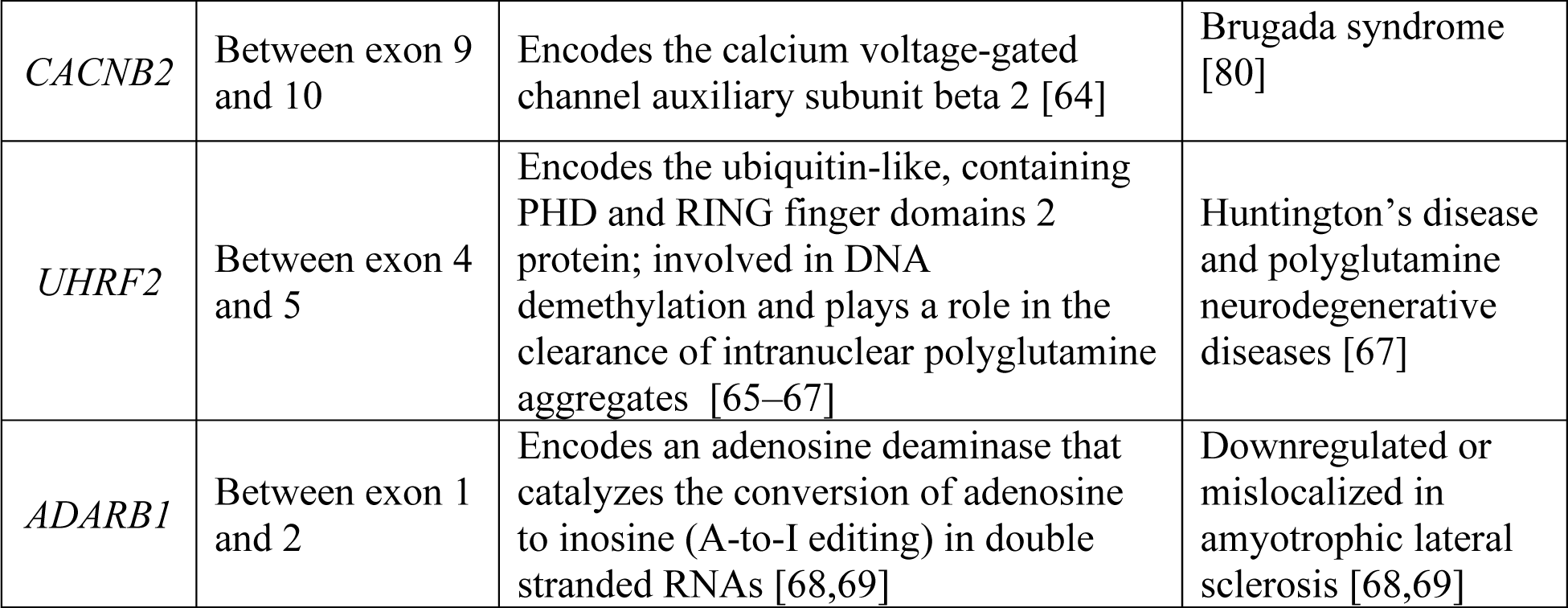
Targets with high fold change of cryptic exon inclusion upon loss of MATR3.

Next, we determined whether MATR3 is responsible for the repression of these cryptic exons by testing whether exogenous MATR3 can rescue cryptic exon inclusion defects upon knockdown of endogenous MATR3 in HeLa cells. Exogenous MATR3 was introduced by transfecting HeLa cells with pcDNA-FLAG-HA-MATR3, encoding only the coding region of human *MATR3.* To avoid knocking down the exogenous *MATR3* mRNA, we used siRNA targeting the 3ʹ untranslated region (UTR) of *MATR3* mRNA and confirmed its knockdown efficiency using MATR3 primers that bind the 5ʹ UTR (Figure 2A, Supplementary Figure S8A and B). *UQCRC2* cryptic exon inclusion levels were examined in HeLa cells co-transfected with the wildtype (WT) MATR3 construct and MATR3 siRNA or control siRNA using RT-PCR. We found basal levels of cryptic exon inclusion in the control samples, which were further reduced upon MATR3 overexpression (Figure 2A and B). Knockdown of MATR3 significantly increased the inclusion of the cryptic exon, and this phenotype was significantly rescued upon overexpression of exogenous MATR3 (Figure 2A-D). To determine whether cryptic exon inclusion in the other top targets occurs due to loss of MATR3, we evaluated rescue of cryptic exon inclusion and found that exogeneous expression of MATR3 can rescue cryptic exon repression in *PHF20* (Supplementary Figure S9A-B), *CACNB2* (Supplementary Figure S9C-D), *UHRF2* (Figure 2H-I, Supplementary Figure S9E-F), and *ADARB1* (Supplementary Figure S10B-C). Together, these results demonstrate that cryptic exon inclusion in these targets is due to the loss of MATR3, confirming that these cryptic splicing events are regulated by MATR3.

**Figure 2.**
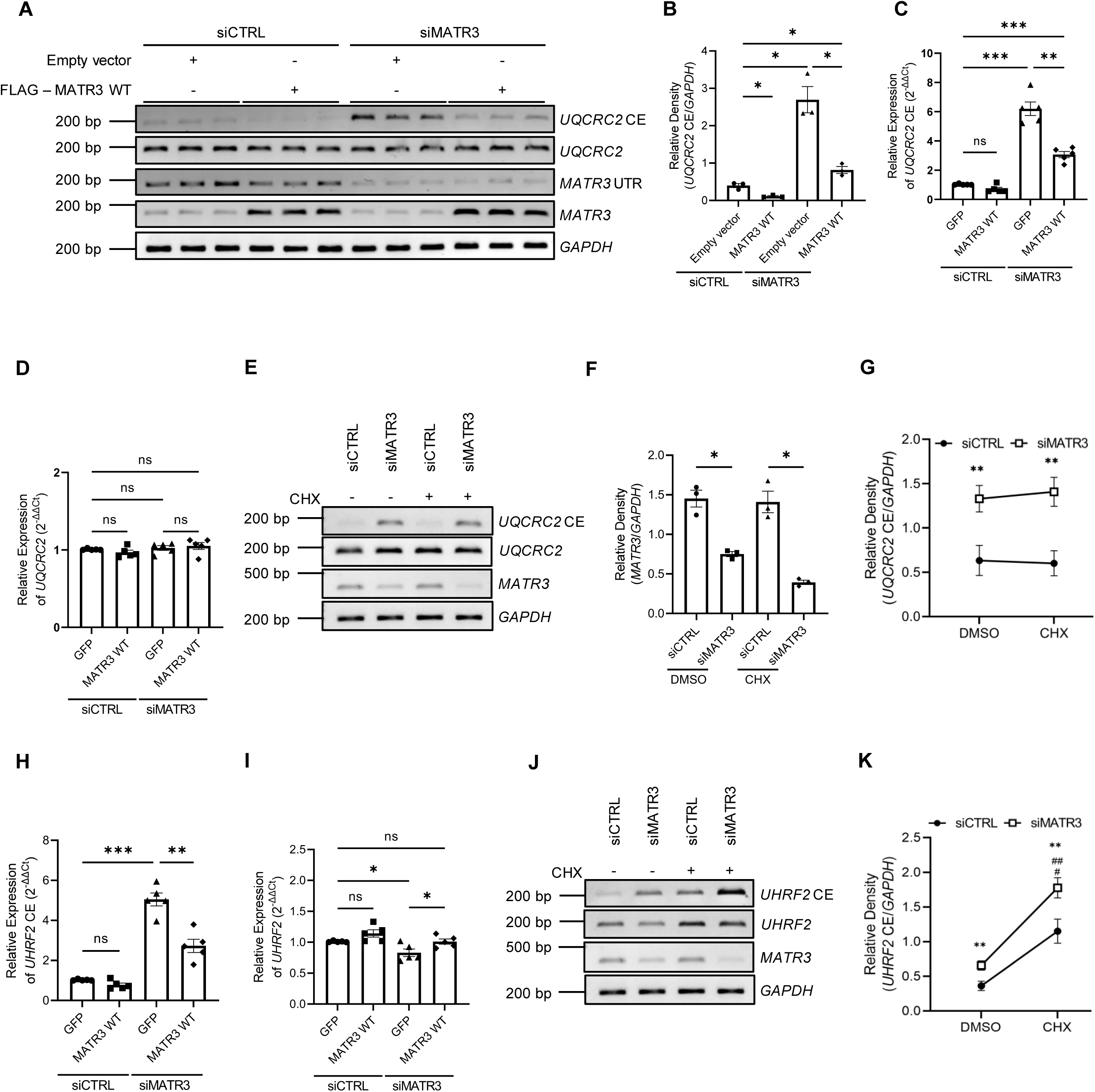
Loss of MATR3 leads to cryptic exon inclusion in *UQCRC2* and *UHRF2*. **A.** Representative RT-PCR gels showing transcript levels of *GAPDH*, *MATR3* UTR and coding region and *UQCRC2* cryptic exon (CE) from total RNA extracted from HeLa cells doubly transfected with nontargeting control siRNA (siCTRL) or siRNA targetingMATR3 (siMATR3) and control vector or FLAG-MATR3 wildtype (WT). **B.** Quantification of cryptic exon inclusion in *UQCRC2* normalized to *GAPDH* (n=3, bar heights depict mean ± SEM, with each datapoint representing a biological replicate, significance determined by Welch’s t-test, * p ≤ 0.05). **C-D.** Relative gene expression (ΔΔC_t_ analysis) of CE-containing *UQCRC2* transcripts or total *UQCRC2* mRNA in HeLa cells doubly transfected with either siCTRL or siMATR3 and FLAG-GFP or FLAG-MATR3 WT as measured by quantitative RT-PCR (n=5, bar heights depict mean ± SEM, with each datapoint representing a biological replicate, significance determined by Welch’s t-test, ns *p* > 0.05, ** *p* ≤ 0.01, *** *p* ≤ 0.001). **E.** Representative gels showing RT-PCR of *UQCRC2 CE, UQCRC2, MATR3,* and *GAPDH* from total RNA extracted from HeLa cells transfected with siRNA and then treated with cycloheximide (CHX) or DMSO control. **F.** Quantification of *MATR3* transcript levels normalized to *GAPDH* to confirm knockdown (n=3, bar heights depict mean ± SEM, with each datapoint representing a biological replicate, significance determined by Welch’s t-test, * p ≤ 0.05). **G.** Quantification of the accumulation of *UQCRC2* transcripts containing the cryptic exon normalized to *GAPDH* (n=3, mean ± SEM are plotted, with each datapoint representing a biological replicate, significance determined by two-way ANOVA, ** p ≤ 0.01 representing the statistical comparison between siCTRL and siMATR3, statistical comparison between DMSO and CHX treatments within each siRNA is not significant). **H-I.** Relative gene expression (ΔΔC_t_ analysis) of CE-containing *UHRF2* transcripts or total *UHRF2* mRNA in HeLa cells doubly transfected with either siCTRL or siMATR3 and FLAG-GFP or FLAG-MATR3 WT as measured by quantitative RT-PCR (n=5, bar heights depict mean ± SEM, with each datapoint representing a biological replicate, significance determined by Welch’s t-test, ns *p* > 0.05, * p ≤ 0.05, ** *p* ≤ 0.01, *** *p* ≤ 0.001). **J.** Representative gels showing RT-PCR of *UHRF2 CE, UHRF2, MATR3,* and *GAPDH* from total RNA extracted from HeLa cells transfected with siRNA and then treated with CHX or DMSO control. **K.** Quantification of the accumulation of *UHRF2* transcripts containing the cryptic exon normalized to *GAPDH* (n=3, mean ± SEM are plotted, with each datapoint representing a biological replicate, significance determined by two-way ANOVA, ** p ≤ 0.01 representing the statistical comparison between siCTRL and siMATR3, ^###^ *p* ≤ 0.001 representing the statistical comparison between DMSO and CHX treatments within each siRNA).

We then examined the downstream consequences of cryptic exon inclusion. Previously, inclusion of cryptic exons has been shown to induce nonsense mediated decay (NMD) of the transcript through the introduction of a premature termination codon (PTC) [5]. Of the cryptic splicing events that we identified upon MATR3 knockdown, we detected 165 cryptic junctions with NMD potential (Supplementary Data File 3). Specifically, we found PTCs in *UHRF2*, *CACNB2* and *ADARB1* following cryptic exon inclusion. To validate this, we used an *in vitro* cycloheximide sensitivity assay [70]. Cells were transfected with siRNA targeting MATR3 and treated with cycloheximide, an indirect inhibitor of NMD, and the accumulation of transcripts containing the cryptic exon was assessed using RT-PCR (Figure 2E-G and 2J-K). We validated that *UHRF2* transcripts containing the cryptic exon are likely targeted for NMD as treatment with cycloheximide led to a significant accumulation of *UHRF2* transcripts containing the cryptic exon (Figure 2J-K). In contrast, we found that *UQCRC2* transcripts containing the cryptic exon were not sensitive to cycloheximide treatment (Figure 2E-G), suggesting they are not targeted for NMD.

We next investigated which protein domains in MATR3 are required for its cryptic splicing repression function (Figure 3A). We cloned mutants of MATR3 lacking RRM1, RRM2 or both RRM domains into the pcDNA-FLAG-HA vector and evaluated the splicing ability of the MATR3 RRM domain deletion (Δ) mutants on the cryptic exon in *UQCRC2* mRNA. After confirming their expression (Supplementary Figure S11A), we co-transfected HeLa cells with either MATR3 WT or the domain deletion (Δ) mutant constructs with MATR3 siRNA or control siRNA and examined the cryptic exon inclusion levels in *UQCRC2* mRNA by RT-PCR. We found that MATR3 ΔRRM1 significantly rescues the cryptic exon inclusion defects to similar levels as MATR3 WT, whereas MATR3 ΔRRM2 and the double RRM deletion mutant do not (Figure 3B and C). This result demonstrates that the RRM2 domain is required for the cryptic splicing repression ability of MATR3. We also examined MATR3 mutants that lack ZF1, ZF2 or both ZF domains. After confirming their expression (Supplementary Figure S11B), we found that all three ZF deletion mutants are able to significantly rescue the cryptic exon inclusion defects to similar levels as MATR3 WT (Figure 3D and E), thus demonstrating that neither ZF domain is required for the cryptic splicing repression ability of MATR3.

**Figure 3.**
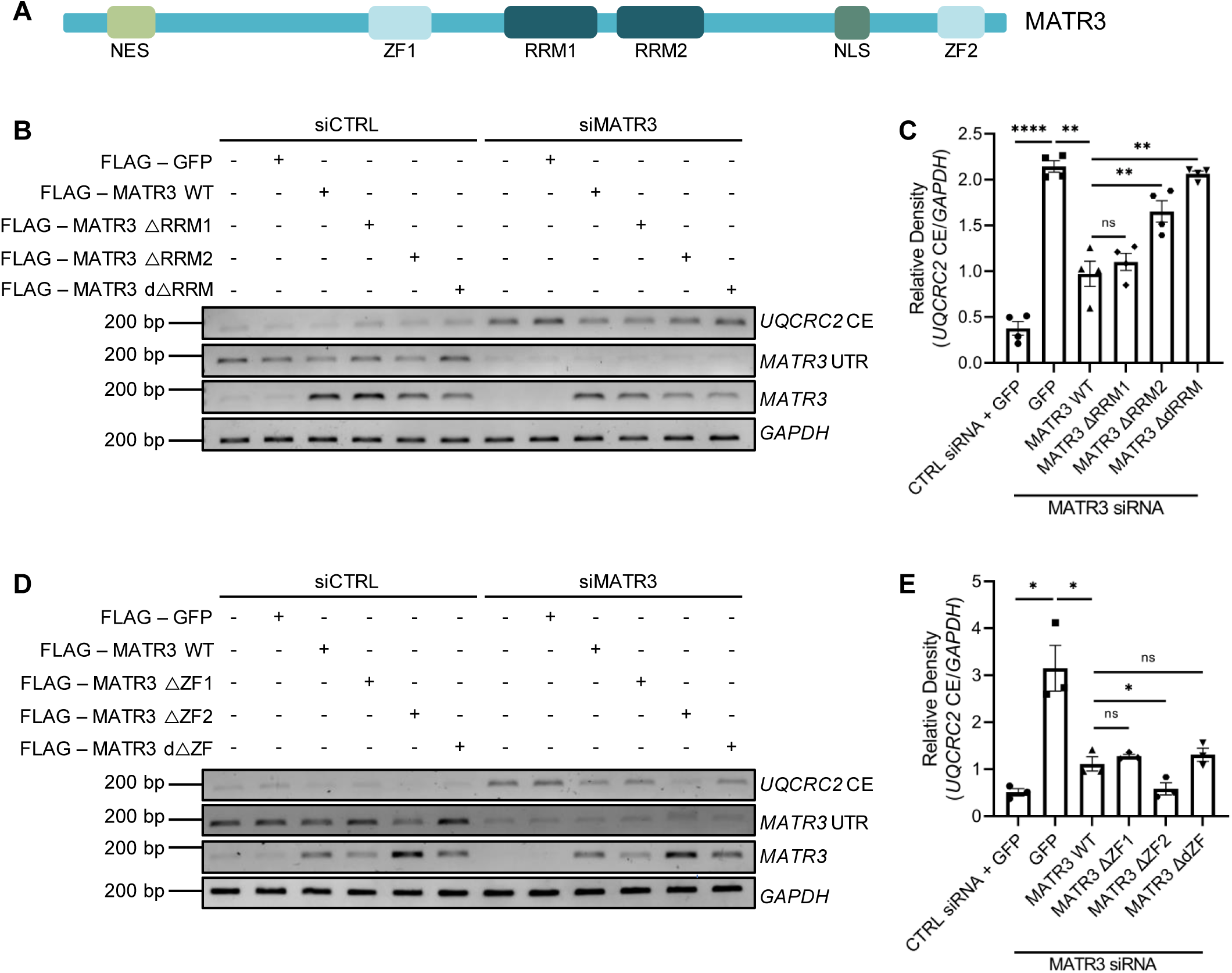
RT-PCR analysis shows that the RRM2 domain is required for MATR3 cryptic splicing repression in *UQCRC2*. **A.** Schematic representation of MATR3 domains showing nuclear export signal (NES), zinc finger domains (ZF), RNA recognition motifs (RRM) and nuclear localization signal (NLS). **B.** Representative gels showing RT-PCR for *GAPDH*, *MATR3* UTR and coding region, and *UQCRC2* cryptic exon (CE) from total RNA extracted from HeLa cells transfected with siRNA and MATR3 vectors. **C.** Quantification of *UQCRC2* cryptic exon inclusion normalized to *GAPDH* (n=4, bar heights depict mean ± SEM, with each datapoint representing a biological replicate, with a total of 4 independent experiments performed, significance determined by Welch’s t-test, * p ≤ 0.05, ** p ≤ 0.01, **** p ≤ 0.0001 ns=not significant). **D.** Representative gels showing RT-PCR for *GAPDH*, *MATR3* UTR and coding region and *UQCRC2* cryptic exon (CE) from total RNA extracted from HeLa cells transfected with siRNA and MATR3 vectors. **E.** Quantification of *UQCRC2* cryptic exon inclusion normalized to *GAPDH* (n=3, bar heights depict mean ± SEM, with each datapoint representing a biological replicate, with a total of 3 independent experiments performed, significance determined by Welch’s t-test, * p ≤ 0.05, ns=not significant).

To determine whether MATR3 regulates cryptic splicing through the binding of its targets, we performed RNA immunoprecipitation to detect the pulldown of target transcripts by MATR3 immune complexes. As previous studies have shown that MATR3 binds RNA predominantly via the RRM2 domain [23], and as we have demonstrated the importance of the RRM2 domain for the cryptic splicing repression function of MATR3, we used MATR3 ΔRRM2 as a negative control. Using a FLAG antibody, we pulled down the FLAG-tagged MATR3, extracted RNA from the immune complexes and performed RT-PCR to evaluate target binding. We assessed the pulldown of the full-length MATR3 protein and the RRM2 deletion mutant protein (Figure 4A), and then evaluated the binding of *UQCRC2* and *CACNB2* mRNAs to the immune complexes by calculating the binding ratio of these target transcripts to the protein pulldown. We found that MATR3 WT readily binds both targets, while MATR3 ΔRRM2 binds with a significantly reduced affinity (Figure 4B-D). These results demonstrate that both *UQCRC2* and *CACNB2* mRNAs are bound by MATR3-associated complexes, and that this binding is mediated by the RRM2 domain.

**Figure 4.**
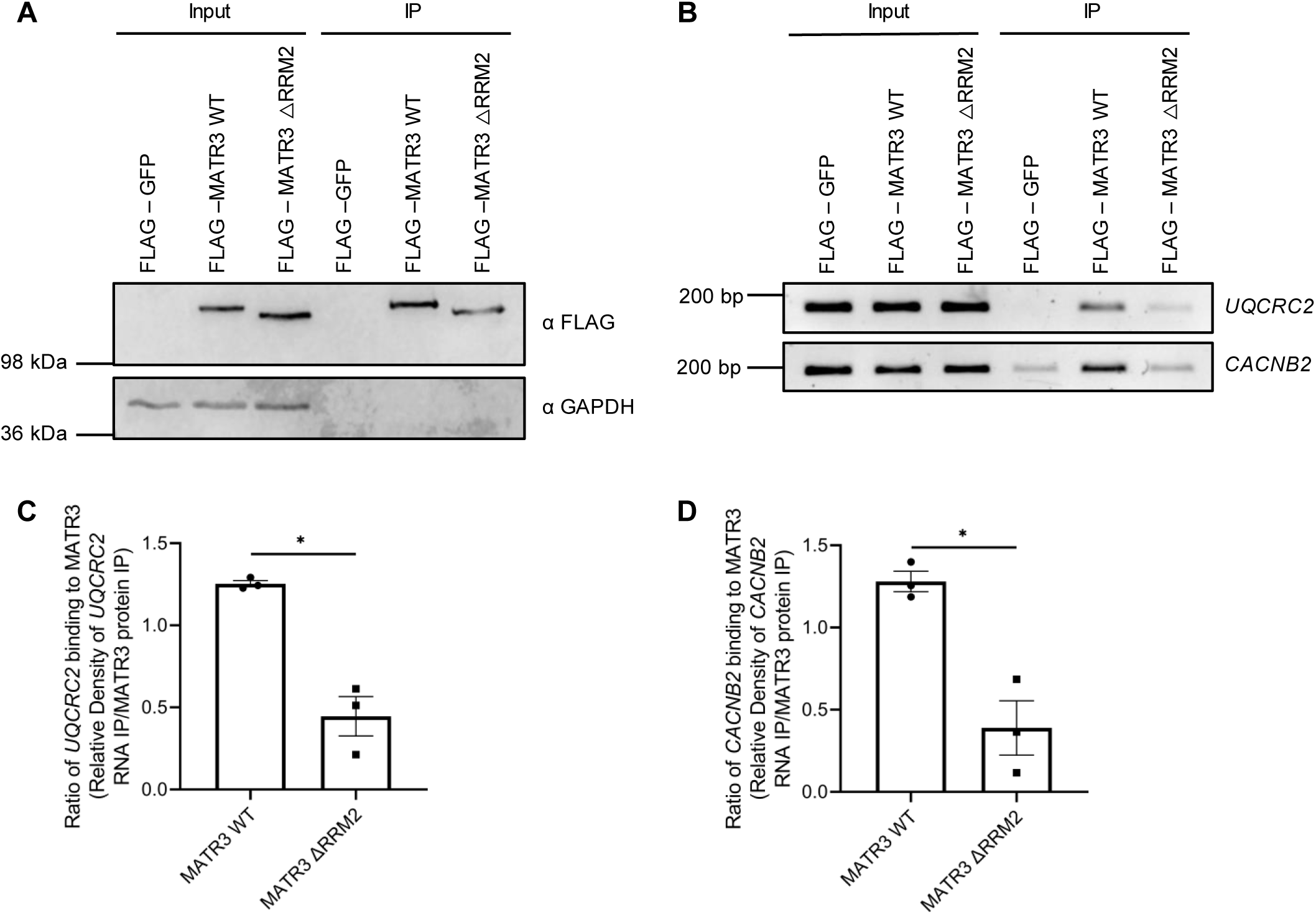
RNA immunoprecipitation analysis shows enrichment of *UQCRC2* and *CACNB2* mRNA in MATR3 WT pulldown. **A.** Western blot showing immunoprecipitation (IP) of MATR3 WT and MATR3 ΔRRM2. **B.** RT-PCR of *UQCRC2* and *CACNB2* from cDNA of input and IP samples. **C.** Quantification of binding ratio of *UQCRC2* to MATR3 (n=3, bar heights depict mean ± SEM, with each datapoint representing a biological replicate, with a total of 3 independent experiments performed, significance determined by Welch’s t-test, * p ≤ 0.05). **D.** Quantification of binding ratio of *CACNB2* to MATR3 (n=3, bar heights depict mean ± SEM, with each datapoint representing a biological replicate, with a total of 3 independent experiments performed, significance determined by Welch’s t-test, * p ≤ 0.05).

Next, we evaluated the effect of an ALS-linked pathogenic variant on MATR3’s cryptic splicing repression function. The S85C pathogenic variant is the most common ALS-linked variant in *MATR3* (Figure 5A). As reported previously [40,71,72], we found that the S85C pathogenic variant lowers the solubility of MATR3 in HeLa cells (Supplementary Figure S12 A-D), therefore decreasing the functional levels of MATR3. We evaluated the effect of MATR3 S85C on cryptic exon inclusion in *UQCRC2* mRNA and found that the overexpression of MATR3 S85C is not able to rescue the cryptic splicing defects upon loss of MATR3 to the same degree as MATR3 WT (Figure 5B and C). To determine whether the effect of this pathogenic variant can be recapitulated in a clinically relevant context, we used iPSC derived neurons harboring the disease-causing S85C pathogenic variant and compared cryptic exon inclusion to the isogenic control. As in HeLa cells, we found decreased soluble MATR3 (Supplementary Figure S12E and F), as well as increased cryptic exon inclusion in *UQCRC2* mRNA in *MATR3^S85C/+^* neurons compared to *MATR3^+/+^* neurons (Figure 5D and E). Together, these results corroborate that the S85C pathogenic variant leads to loss of MATR3 function and that the cryptic exon in *UQCRC2* mRNA is a MATR3 target. We further evaluated the effect of the S85C pathogenic variant on MATR3 binding to target mRNAs using RNA immunoprecipitation. After confirming the immunoprecipitation of MATR3 WT and mutant proteins (Figure 5F), we examined the levels of *UQCRC2* and *CACNB2* mRNA bound to the immune complexes. Although there was less pulldown of MATR3 S85C, possibly due to reduced solubility, we found that MATR3 S85C binds target mRNAs as strongly as MATR3 WT (Figure 5G-I). These results suggest that the S85C pathogenic variant does not affect the RNA binding ability of MATR3.

**Figure 5.**
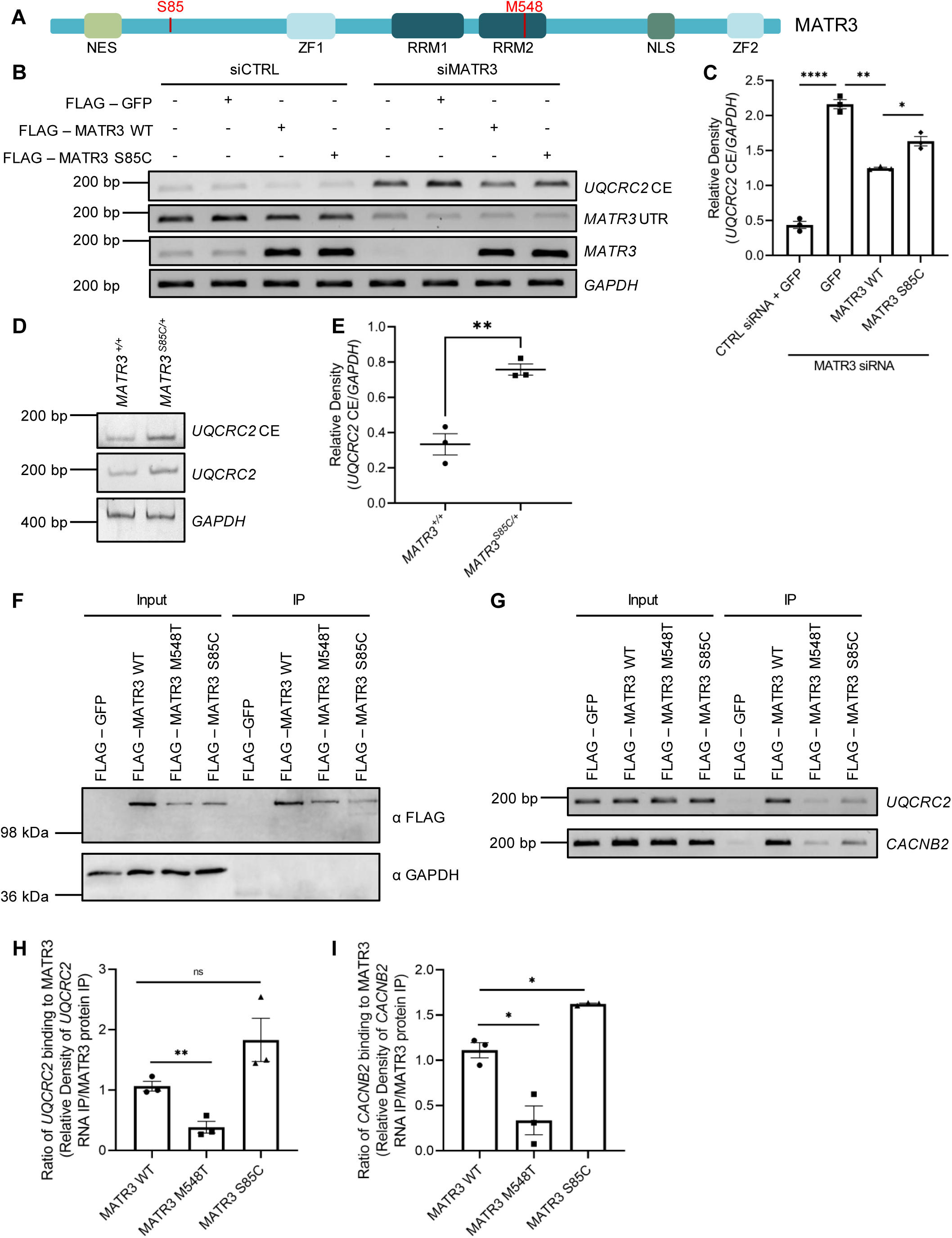
The S85C pathogenic variant affects the cryptic splicing repression function of MATR3 but does not affect its RNA binding, unlike MATR3 M548T. **A.** Schematic representation of MATR3 showing nuclear export signal (NES), zinc finger domains (ZF), RNA recognition motifs (RRM), nuclear localization signal (NLS) and location of the S85C and M548T variants. **B.** Representative gels showing RT-PCR for *GAPDH*, *MATR3* UTR and coding region, and *UQCRC2* cryptic exon (CE) from total RNA extracted from HeLa cells transfected with siRNA and MATR3 vectors. **C.** Quantification of cryptic exon inclusion in *UQCRC2* normalized to *GAPDH* (n=3, bar graph heights depict mean ± SEM, with each datapoint representing a biological replicate, with a total of 3 independent experiments performed, significance determined by Welch’s t-test, * p ≤ 0.05, ** p ≤ 0.01, **** p ≤ 0.0001). **D.** Representative gels showing RT-PCR for *GAPDH* and *UQCRC2* internal and cryptic exon mRNAs from differentiated *MATR3 ^S85C/+^* and isogenic Kolf2.1 control neurons. **E**. Quantitative graph showing an increase in *UQCRC2* cryptic exon mRNA in *MATR3 ^S85C/+^* neurons as compared to control. *GAPDH* mRNA levels were used as a normalization control (n=3, significance determined by Welch’s t-test, ** p ≤ 0.001). **F.** Western blot showing immunoprecipitation (IP) of MATR3 WT, MATR3 M548T, and MATR3 S85C. **G.** RT-PCR of *UQCRC2* and *CACNB2* from cDNA of input and IP samples. **H.** Quantification of binding ratio of *UQCRC2* to MATR3 (n=3, bar graph heights depict mean ± SEM, with each datapoint representing a biological replicate, with a total of 3 independent experiments performed, significance determined by Welch’s t-test, ** p ≤ 0.01, ns=not significant). **I.** Quantification of binding ratio of *CACNB2* to MATR3 (n=3, bar graph heights depict mean ± SEM, with each datapoint representing a biological replicate, with a total of 3 independent experiments performed, significance determined by Welch’s t-test, * p ≤ 0.05).

Recently, a heterozygous *de novo* missense variant in *MATR3* (E436K) was identified in a child with severe neurodegeneration [42]. Examination of protein levels from patient skin fibroblasts revealed a reduction of nuclear MATR3 levels compared to healthy controls, suggesting loss of functional protein levels as a molecular mechanism of disease. This prompted us to look for additional *MATR3* variants associated with early-onset brain disease. Through review of a clinical exome sequencing database, we identified a patient with a heterozygous M548T mutation in *MATR3* exhibiting neurodevelopmental defects including delayed motor milestones, developmental regression and intellectual disability, as well as mitochondrial defects including complex III deficiency (Table 2). Interestingly, the novel MATR3 M548T variant lies within the RRM2 domain (Figure 5A). To investigate how the M548T variant impairs MATR3 function, we generated the MATR3 M548T construct through mutagenesis. We first verified protein expression in HeLa cells and examined the variant’s impact on protein solubility. We found that MATR3 M548T is less soluble compared to MATR3 WT (Figure 6A and C) and correspondingly more insoluble than the WT protein (Figure 6B and D). This result suggests that the M548T variant may cause severe consequences through decreasing functional MATR3 protein levels.

**Figure 6.**
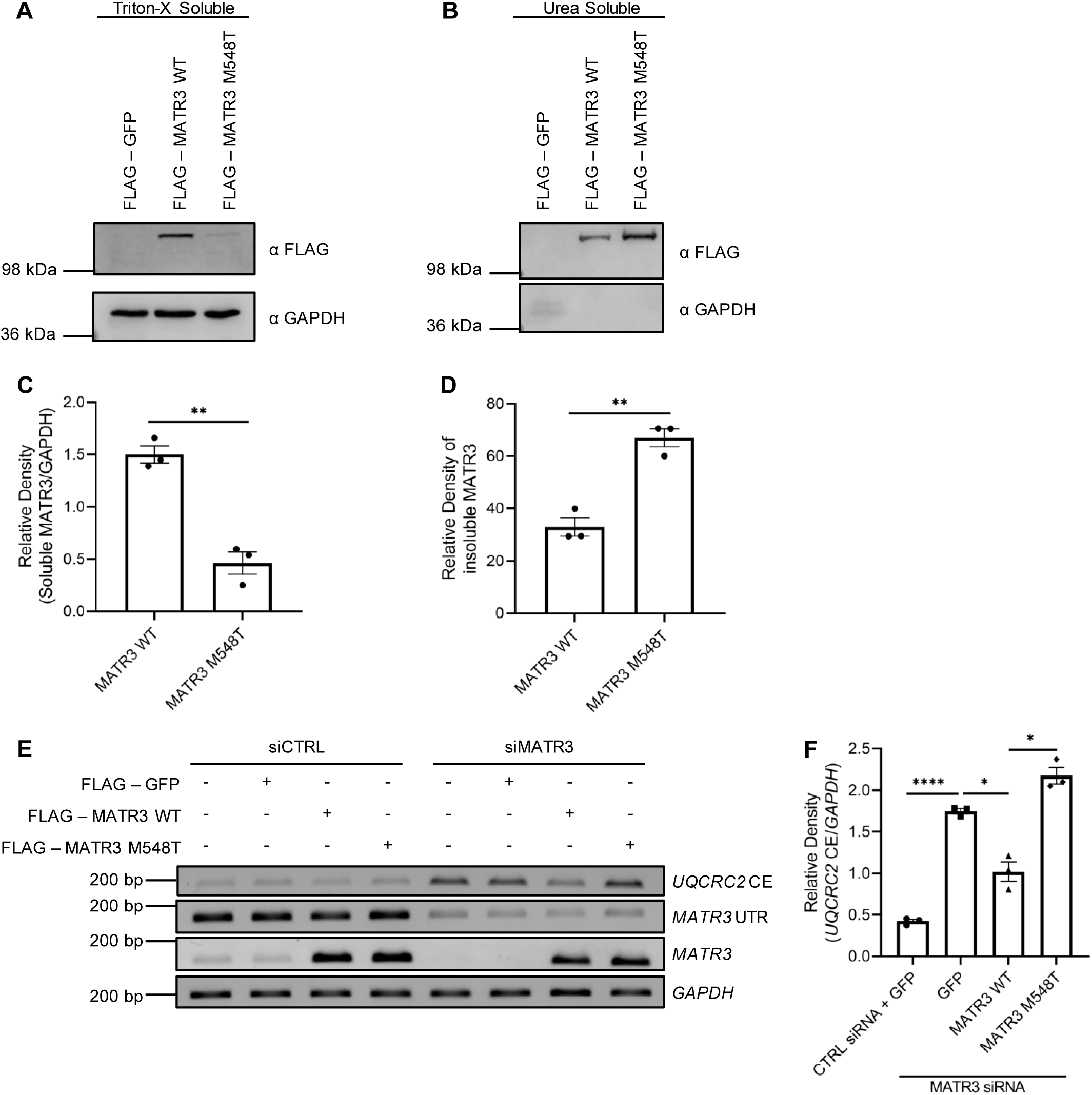
MATR3 M548T is less soluble than MATR3 WT and does not rescue cryptic exon inclusion in *UQCRC2* upon loss of MATR3. **A.** Western blot showing Triton-X soluble (soluble fraction) protein levels of MATR3 WT and MATR3 M548T. **B.** Western blot showing urea soluble (insoluble fraction) protein levels of MATR3 WT and MATR3 M548T. **C.** Quantification of soluble MATR3 levels normalized to GAPDH (n=3, bar heights depict mean ± SEM, with each datapoint representing a biological replicate, with a total of 3 independent experiments performed, significance determined by Welch’s t-test, ** p ≤ 0.01). **D.** Quantification of insoluble MATR3 levels (n=3, bar heights depict mean ± SEM, with each datapoint representing a biological replicate, a total of 3 independent experiments was performed, significance determined by Welch’s t-test, ** p ≤ 0.01). **E.** Representative gels showing RT-PCR for *GAPDH*, *MATR3* UTR and coding region, and *UQCRC2* cryptic exon (CE) from total RNA extracted from HeLa cells transfected with siRNA and MATR3 vectors. **F.** Quantification of cryptic exon inclusion in *UQCRC2* normalized to *GAPDH* (n=3, bar graph heights depict mean ± SEM, with each datapoint representing a biological replicate, with a total of 3 independent experiments performed, significance determined by Welch’s t-test, * p ≤ 0.05, **** p ≤ 0.0001).

**Table 2.** Overview of MATR3 M548T.

| AA Change | BP Change | Location of Variant | Phenotype |
| --- | --- | --- | --- |
| M548T | NM_199189.2: c.1643-T>C | RRM2 domain | Delayed motor milestones, developmental regression, intellectual disability, hypotonia, dysmorphic features (facial droop), muscle pain and fatigue, hypothyroidism, malignant hyperthermia, and elevated creatine kinase.<br>Previous muscle biopsy demonstrated abnormal function of mitochondrial complexes (complex I and complex III deficiency) with mitochondrial myopathy. |

**Table 3.**
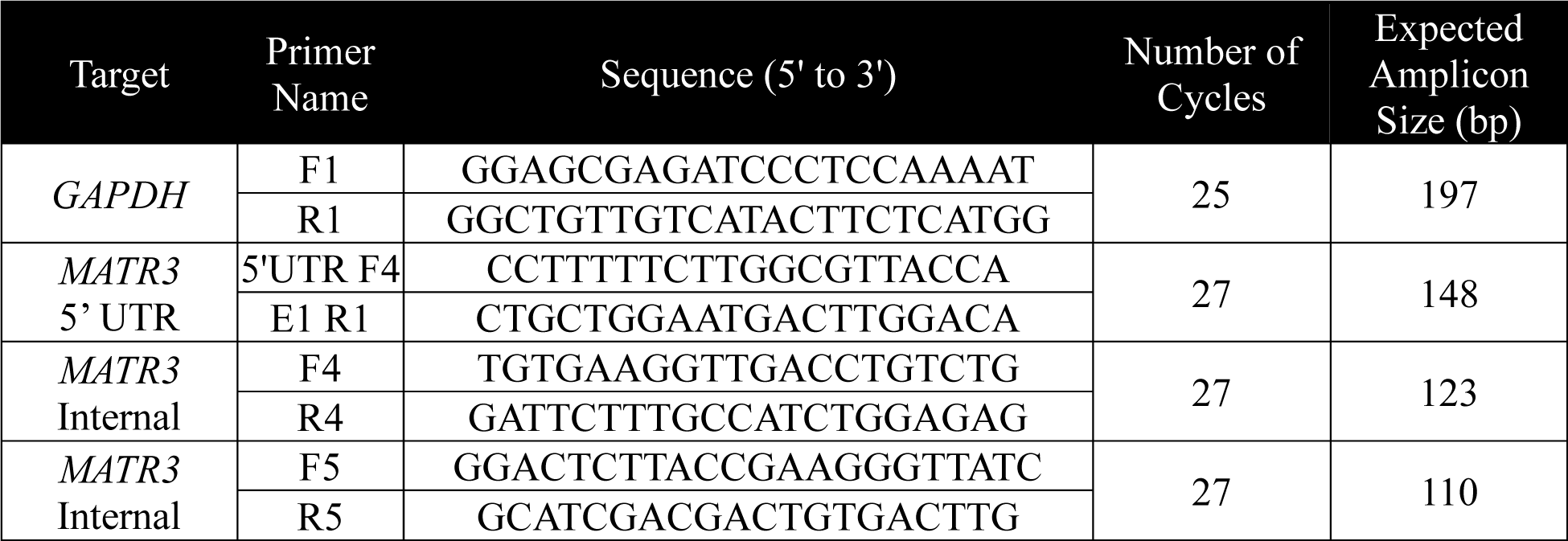
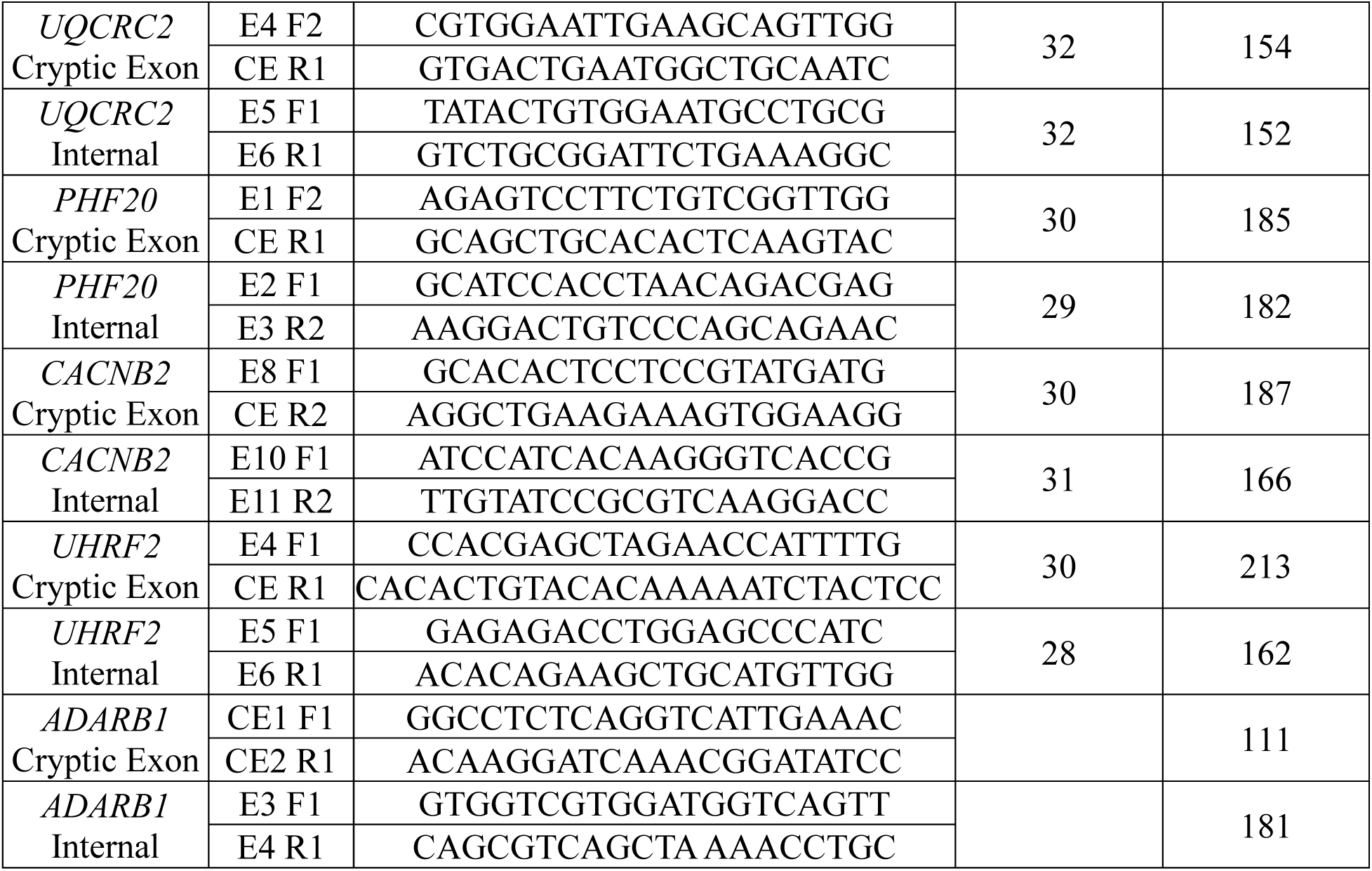
Primers used for RT-PCR. E refers to the exon number, CE refers to the cryptic exon.

Furthermore, we evaluated the functional effect of the MATR3 M548T variant on cryptic exon inclusion in *UQCRC2* mRNA. We found that MATR3 M548T is not able to rescue the cryptic exon inclusion in *UQCRC2* mRNA upon loss of MATR3 compared to MATR3 WT (Figure 6E and F). In addition, we found that MATR3 M548T binds the mRNA targets less strongly than the WT form (Figure 5F-I), suggesting that this variant may also disrupt the RNA binding ability of the RRM2 domain. Together, these results suggest that the failure of MATR3 M548T to rescue cryptic exon inclusion to WT levels may be driven by both the decreased functional protein levels and impaired RNA binding ability.

## Conclusion

In this study, we investigated the role of MATR3 in cryptic splicing within functional genes and the impact of disease-associated variants on its cryptic splicing repression ability in human cell lines. Through rescue experiments with exogenous MATR3, we determined that increased cryptic exon inclusion in *UQCRC2*, *PHF20*, *CACNB2, UHRF2*, and *ADARB1* mRNA is due to the loss of MATR3, indicating that MATR3 is responsible for cryptic exon repression in these targets. We also evaluated the functional effect of domain mutants of MATR3 and found that the RRM2 domain is required for the cryptic splicing repression function of MATR3. Through RNA immunoprecipitation, we demonstrated that *UQCRC2* and *CACNB2* mRNA are bound by MATR3 immune complexes, and that this binding is abolished when the RRM2 domain is lost. Together these results suggest that the RRM2 domain is required for both the cryptic splicing repression function of MATR3 and the binding of target transcripts. These data are supported by previous studies that have shown that the RRM2 domain has RNA binding ability and is required to bind several small noncoding RNAs and mRNAs [18,23,24]. Furthermore, our results reveal that two disease-associated variants differentially affect MATR3 properties and/or function, which may help explain the differing phenotypes associated with the variants. The adult-onset ALS-linked S85C pathogenic variant lies in the N-terminal disordered region, while the neurodevelopmental disease-associated M548T variant lies in the RRM2 domain (Figure 5A). We found that both variants lead to loss of MATR3 function and impair cryptic splicing repression; however, the two variants appear to affect MATR3 function to a different degree. The S85C pathogenic variant increases cryptic exon inclusion, possibly through reducing soluble MATR3 protein levels, while the M548T variant may impact the cryptic splicing repression function of MATR3, possibly by affecting both soluble protein levels and RNA binding ability, therefore, possibly explaining the increased severity of consequences and earlier symptom onset.

We show that extensive cryptic splicing changes occur upon loss of MATR3, and we have identified MATR3-regulated splicing events in *UQCRC2*, *PHF20, CACNB2*, *UHRF2,* and *ADARB1* mRNA. We also found increased pyrimidine-rich sequences upstream of cryptic junctions and increased MATR3 binding near the cryptic exons, in accordance with previous studies [24,25]. These results suggest that these targets may be direct targets of MATR3. Differential gene expression data from DESeq2 and mRNA levels from RT-PCR showed a decrease in *PHF20*, *UHRF2,* and *ADARB1* levels upon cryptic exon inclusion, while the levels of *UQCRC2* and *CACNB2* showed no significant change. Previously, inclusion of cryptic exons has been shown to induce NMD of the transcript through the introduction of a PTC, resulting in decreased protein expression [5]. Cryptic exon inclusion in*UHRF2, CACNB2,* and *ADARB1* are predicted to introduce a PTC. We have validated *UHRF2* transcripts containing the cryptic exon to be probable NMD targets, while *UQCRC2* transcripts are not. These results along with the presence of PTC in *ADARB1* corroborate Differential Gene Expression (DEG) analysis.. Further investigation is needed to examine the downstream consequences of cryptic exon inclusion in *PHF20* and *CACNB2*. The cryptic exon in *PHF20* does not encode a PTC but is located in the 5’UTR which is important for transcript stability. Reduction in the stability of *PHF20* transcripts containing the cryptic exon may lead to the downregulation observed in the DEG analysis. Future experiments should evaluate the rate of decay of these target transcripts and examine if the inclusion of the cryptic exon affects the transcriptional regulation, protein translation, localization, protein structure and/or interactor binding of the target. This would provide insights into the downstream consequences of cryptic exon inclusion in the target transcripts.

Another ALS gene, *TARDBP* (TDP-43), is involved in repression of cryptic splicing [8,9,73]. Recent studies showed that loss of TDP-43 leads to cryptic exon inclusion in *UNC13A* mRNA, linking TDP-43 to one of the highest risk factor genes of ALS [14,15]. In addition to *UNC13A*, loss of TDP-43 also leads to cryptic exon inclusion in and downregulation of *STMN2*, a protein that is important for neural growth and repair [74]. Notably, total or partial depletion of STMN2 causes motor deficits in mice which can be rescued by exogenous expression of STMN2 [75,76]. Furthermore, QRL-201, an antisense oligonucleotide designed to correct *STMN2* splicing and restore STMN2 expression, is currently recruiting for a Phase 1 clinical trial (NCT05633459). These discoveries of the role of TDP-43 in cryptic splicing raise the question as to whether other ALS-linked RNA-binding proteins are involved in cryptic splicing regulation and whether the downstream cryptic splicing events are shared, which may explain the underlying ALS pathogenesis [16]. Notably, one recent study identified cryptic exon inclusion in *UQCRC2* mRNA upon knockdown of TDP-43 [14]. Although this event lies downstream of the MATR3 dependent event, this gene could be a possible shared target with MATR3. Further studies are required to answer these questions and identify shared cryptic splicing targets between TDP-43, MATR3 and possibly other RNA-binding proteins.

In this study, we performed the first characterization of the neurodevelopmental disease associated M548T variant in *MATR3*. The M548 residue is located in the second alpha helix of the RRM2 domain [21], and is likely buried and part of the hydrophobic core due to its nonpolar and hydrophobic nature (Supplementary Figure S13). The amino acid substitution to threonine, which is polar, may destabilize the RRM2 domain, resulting in misfolding of the domain. This may lead to the formation of insoluble aggregates, consequently leading to decreased solubility. As RRM domains bind RNA at aromatic residues on the surface of the beta sheets [77], it is unlikely that the M548 residue interacts with the RNA directly. However, misfolding of the RRM2 domain may prevent key residues from contacting RNA, thereby impairing the RNA binding ability of MATR3 M548T. Future experiments are required to investigate how the M548T variant impacts the overall structure of MATR3.

The novel M548T variant in *MATR3* was identified in a patient presenting with neurodevelopmental phenotypes (Table 2) and mitochondrial complex III deficiency. This is particularly interesting as UQCRC2 is a subunit of complex III [62], and variants in *UQCRC2* have been associated with mitochondrial complex III deficiency [78]. This evidence supports the functional link between *UQCRC2* mRNA and MATR3. Future experiments are required to determine how the M548T variant in *MATR3* drives clinical phenotypes by examining the downstream consequences of the variant and its link to *UQCRC2*.

It is interesting to note that the clinical presentation depends on where the variants are located in *MATR3*. Since 2014, more than a dozen ALS-linked missense variants have been identified in *MATR3* with most patients presenting symptoms as adults [16,31,79]. Most of these variants reside outside the functional domains, within disordered regions near the N- or C-termini [16]. Very recently, a *de novo* missense variant E436K in *MATR3* was identified in a child with severe neurodegeneration; interestingly, the E436K variant resides in the RRM1 domain of MATR3 [42]. While inheritance of the M548T variant in our patient has not been determined, the patient’s phenotype, combined with our functional characterization of the variant add support to the hypothesis that disruption of the important functional domains may lead to earlier devastating consequences.

In summary, our findings reveal that MATR3 plays a role in repressing cryptic exon inclusion within functional genes and that the RRM2 domain is the most crucial domain required for this role. Through examination of the effect of two disease-linked variants on the molecular function of MATR3, our findings provide a possible explanation of how different disease-linked variants in *MATR3* may lead to diverse consequences.

## Acknowledgements

We thank the members of the Park laboratory for insightful discussions on this project. We appreciate the support from SickKids TCAG for RNA sequencing. We are grateful to Henrietta Lacks for the HeLa cell line and to her surviving family members for their contributions to biomedical research. This study was supported by Natural Sciences and Engineering Research Council of Canada (NSERC), Canada Research Chairs program and SickKids start-up funds to JP, and United States Department of Agriculture (USDA/ARS) under Cooperative Agreement No. 58-3092-0-001 and Duncan NRI Zoghbi Scholar Award to HKY. MK was supported by the Canada Graduate Scholarship, Ontario Graduate Scholarship, and SickKids Restracomp Scholarship. JRS was supported by a Catalyzing the Talent Pipeline Scholarship from the David Dime Family Catalyst Initiative in Molecular Genetics at the University of Toronto. This paper was generated with contributions from University of Toronto Master of Science theses from Xiao Xiao Lily Chen and Mashiat Khan.

## Author contributions

MK, XXLC, HKY and JP conceptualized and developed methodology for the experiments. MK, XXLC, JRS, SK, JY, RvB, MMMY, and JP performed the experiments and analyzed the data. MD, YW and HKY performed the bioinformatics analysis. QT and JAR provided patient information. ZL, UBP, QT, HKY and JP provided project supervision. MK, XXLC, MD, JRS, HKY and JP wrote the manuscript. All authors revised and approved the manuscript.

## Conflict of interest

The Department of Molecular and Human Genetics at Baylor College of Medicine receives revenue from clinical genetic testing completed at Baylor Genetics Laboratories.

