## Supplementary File for "Differential effects of MATR3 variants on its cryptic splicing repression function"

#### Table of Contents


Supplementary Figure 1. Confirmation of MATR3 knockdown through RT-PCR.

Supplementary Figure 2. RNA sequencing reveals differentially expressed genes upon MATR3 knockdown.

Supplementary Figure 3. Knockdown of MATR3 results in alternative splicing events.

Supplementary Figure 4. Cryptic splicing events upon MATR3 knockdown.

Supplementary Figure 5. Validation of cryptic exon inclusion in *UQCRC2* by RT-PCR.

Supplementary Figure 6. Validation of cryptic exon inclusion in *PHF20* by RT-PCR.

Supplementary Figure 7. Validation of cryptic exon inclusion in *CACNB2* by RT-PCR.

Supplementary Figure 8. Optimization of MATR3 WT rescue experiment.

Supplementary Figure 9. RT-PCR confirms cryptic exon inclusion in *PHF20, CACNB2,* and *UHRF2* upon loss of MATR3 was rescued by MATR3 WT.

Supplementary Figure 10. Cryptic splicing events upon MATR3 knockdown in *ADARB1*

Supplementary Figure 11. Expression of MATR3 domain deletion mutants.

Supplementary Figure 12. Soluble and insoluble fractionation of MATR3 S85C shows it is less soluble than MATR3 WT.

Supplementary Figure 13. NMR structure of the RRM2 domain of MATR3.

**Figure S1**

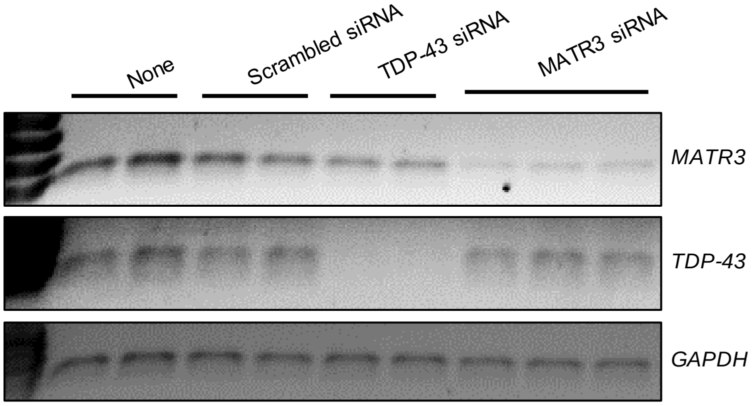


**Supplementary Figure 1. Confirmation of MATR3 knockdown through RT-PCR.** Representative gels showing RT-PCR from total RNA extracted from HeLa cells transfected with scrambled siRNA, TDP-43 siRNA and MATR3 siRNA showing *MATR3*, *TDP-43* and *GAPDH.*

#### Figure S2

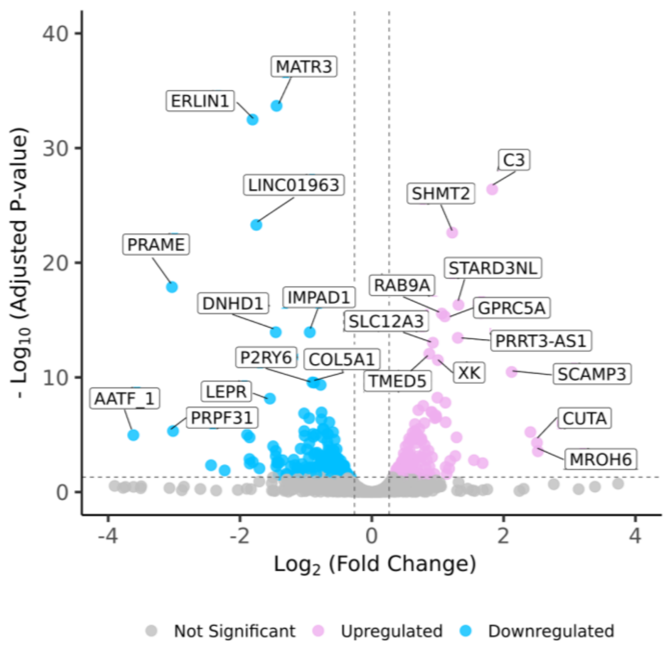


**Supplementary Figure 2. RNA sequencing reveals differentially expressed genes upon MATR3 knockdown.** Volcano plot illustrating genes that are significantly upregulated (pink) and downregulated (blue) upon MATR3 knockdown. The horizontal dotted line indicates the adjusted P-value cutoff of 0.05, and the two vertical dotted lines indicate the $\log_2$ fold change cutoff of 20% (-0.263 or 0.263). A few of the most upregulated and downregulated genes are labeled.

#### Figure S3

| Alternative Splicing Types | | # Splicing Events upon MATR3 KD (% of Total Events) |
|---|---|---|
| Skipped Exon (SE) | 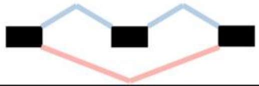 | 1358 (63.7%) |
| Alternative 5’ Splice Site (A5SS) | 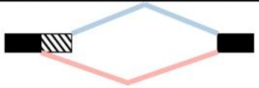 | 229 (10.7%) |
| Alternative 3’ Splice Site (A3SS) | 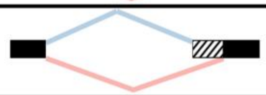 | 272 (12.8%) |
| Mutually Exclusive Exons (MXE) | 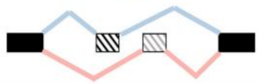 | 152 (7.1%) |
| Retained Intron (RI) | 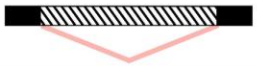 | 120 (5.6%) |
| Total | | 2131 |

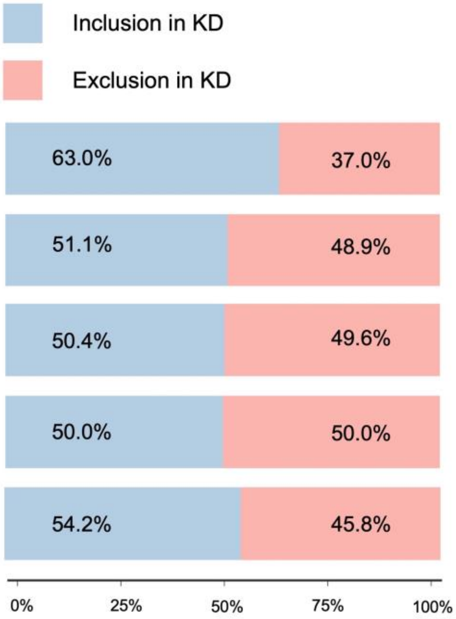


**Supplementary Figure 3. Knockdown of MATR3 results in alternative splicing events.** Alternative splicing events summarized by category: skipped exon (SE), alternative 5’ splice site (A5SS), alternative 3’ splice site (A3SS), mutually exclusive exons (MXE), and retained intron (RI). Significant events identified using a false discovery rate of ≤ 0.05 and an inclusion level difference ≥ 0.2 or ≤ -0.2. The table includes percentages of the significant events per category out of the total significant events. The sidebars illustrate the proportions of inclusion and exclusion events per category. Blue represents events with greater exon inclusion in the knockdown samples, while pink represents events with greater exon exclusion in the knockdown samples. In the illustrations for each category, a blue line indicates an ‘inclusion’ event while the pink line indicates an ‘exclusion’ event.

### Figure S4

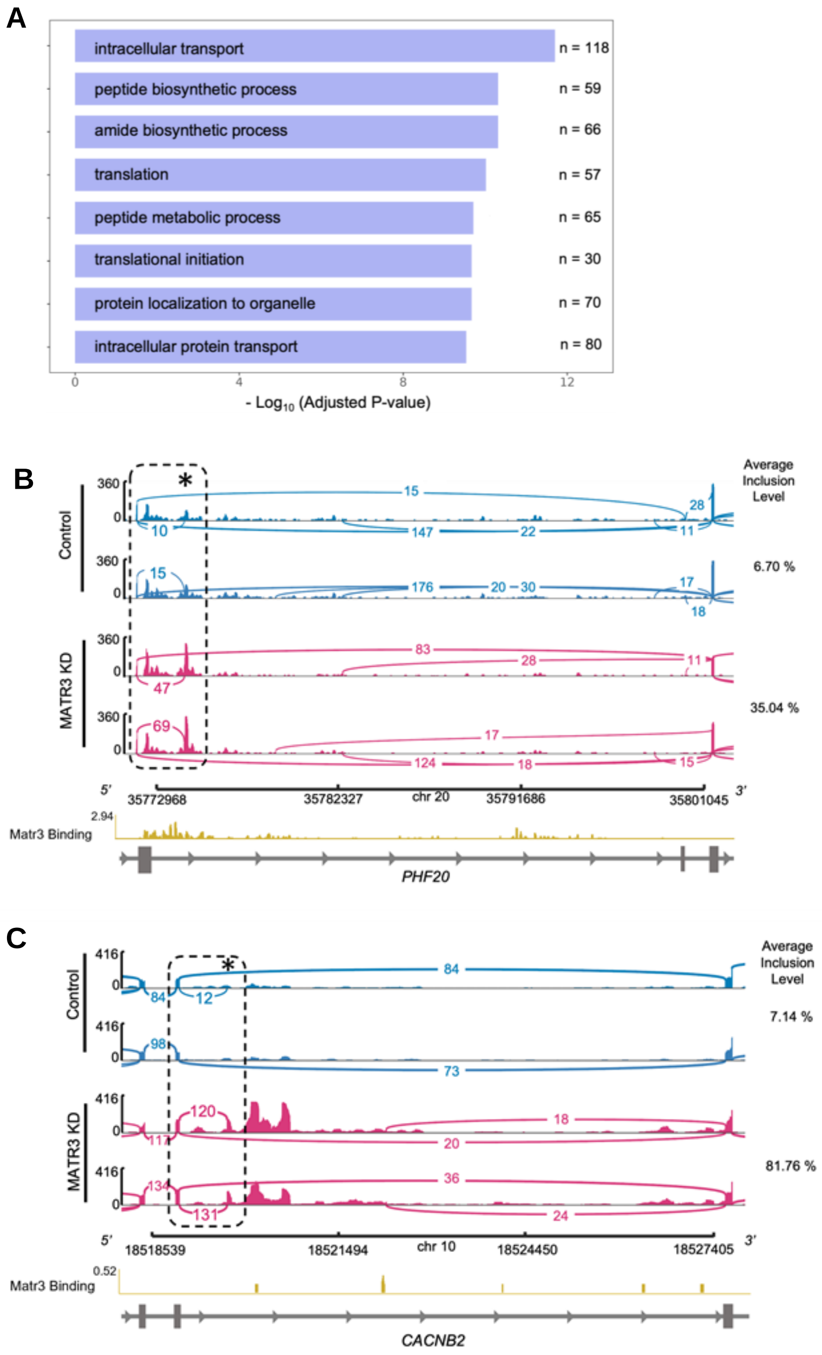


**Supplementary Figure 4. Cryptic splicing events upon MATR3 knockdown. A.** GO term enrichment of the genes containing cryptic splicing events which occurred upon MATR3 knockdown. **B-C.** Sashimi plot generated using IGV genome browser illustrating the read coverage for two biological replicates of control (blue) and *MATR3* knockdown (pink) for a region of *PHF20,* and *CACNB2* transcribed from left to right. The bar graph represents read depth, and the arcs represent the number of reads connecting each splice junction, with junction coverage minimum set to 10. The region within black dashed lines represents the cryptic junction of interest with the cryptic exon marked by an asterisk, and inclusion levels of the cryptic exon within this region are shown as percentages to the right of the tracks. MATR3 binding analyzed from CLIP-Seq data from Coelho *et al*., 2015 is shown in yellow.

#### Figure S5

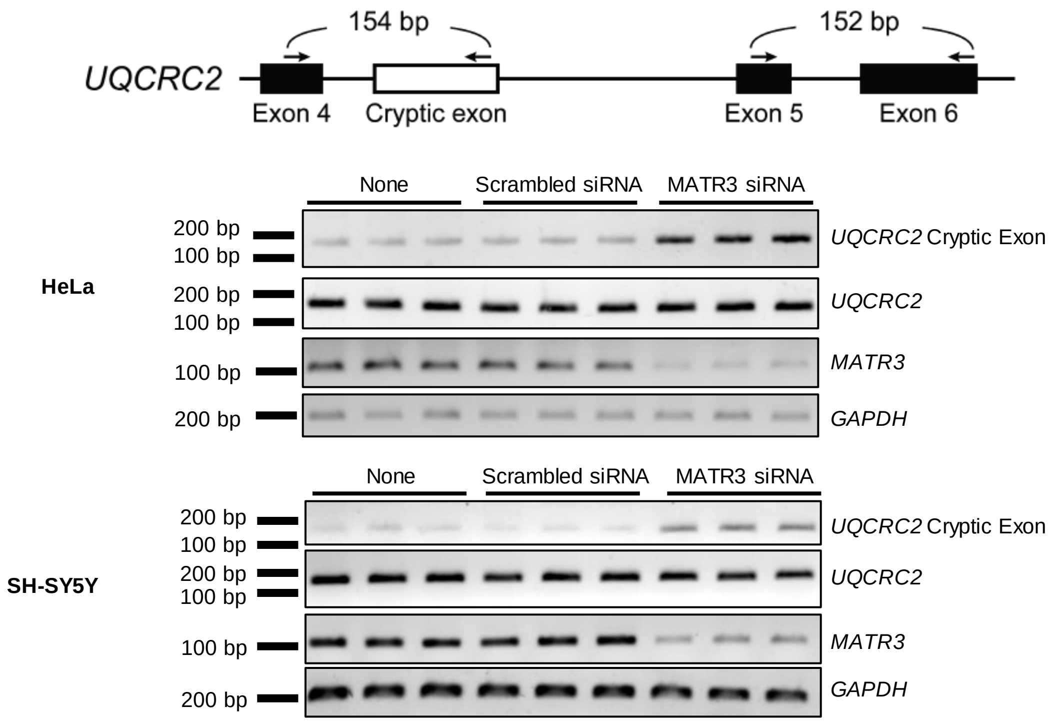


**Supplementary Figure 5. Validation of cryptic exon inclusion in *UQCRC2* by RT-PCR.** The location of the primers used for RT-PCR and the size of the RT-PCR products are denoted in the schematic. RT-PCR in HeLa and SH-SY5Y cells show more cryptic exon inclusion in *UQCRC2* transcripts in all three biological replicates of MATR3 knockdown samples compared. RT-PCR of exon 5 to exon 6 are used as internal controls.

**Figure S6**

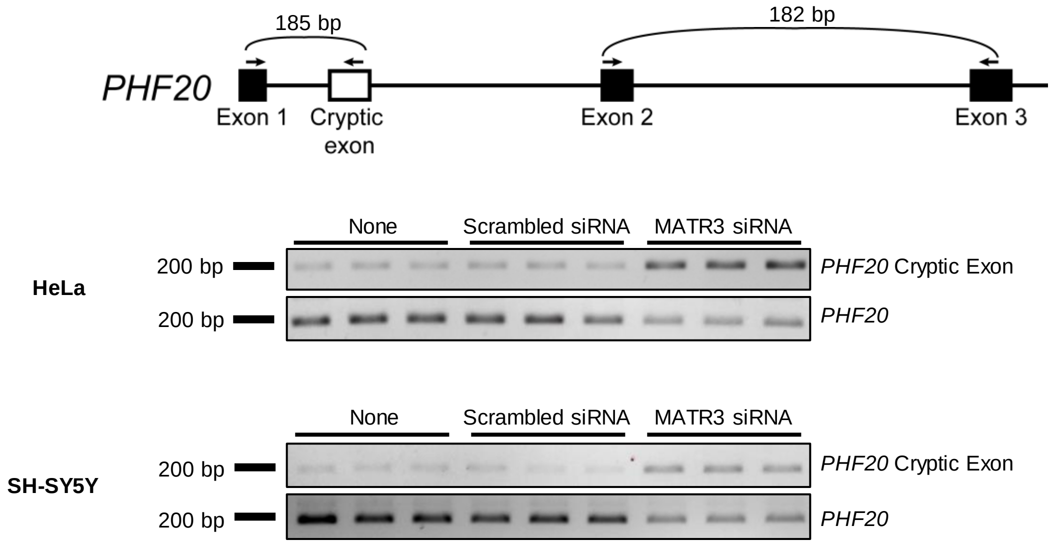


**Supplementary Figure 6. Validation of cryptic exon inclusion in *PHF20*.** The location of the primers used for RT-PCR and the size of the RT-PCR products are denoted in the diagram. RT-PCR in HeLa and SH-SY5Y cells show more cryptic exon inclusion in *PHF20* transcripts in all three replicates of MATR3 KD samples compared to scrambled siRNA and non-transfected controls. RT-PCR of exon 2 to exon 3 are used as internal controls.

#### Figure S7

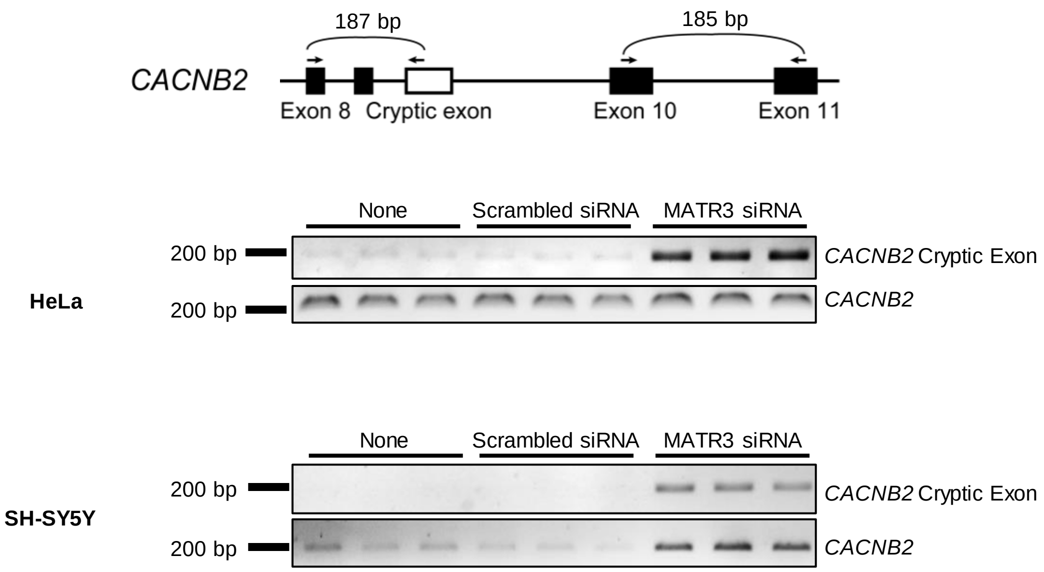


**Supplementary Figure 7. Validation of cryptic exon inclusion in *CACNB2*.** The location of the primers used for RT-PCR and the size of the RT-PCR products are denoted in the diagram. RT-PCR in HeLa and SH-SY5Y cells show more cryptic exon inclusion in *CACNB2* transcripts in all three replicates of MATR3 KD samples compared to scrambled siRNA and non-transfected controls. RT-PCR of exon 10 to exon 11 is used as an internal control.

#### Figure S8

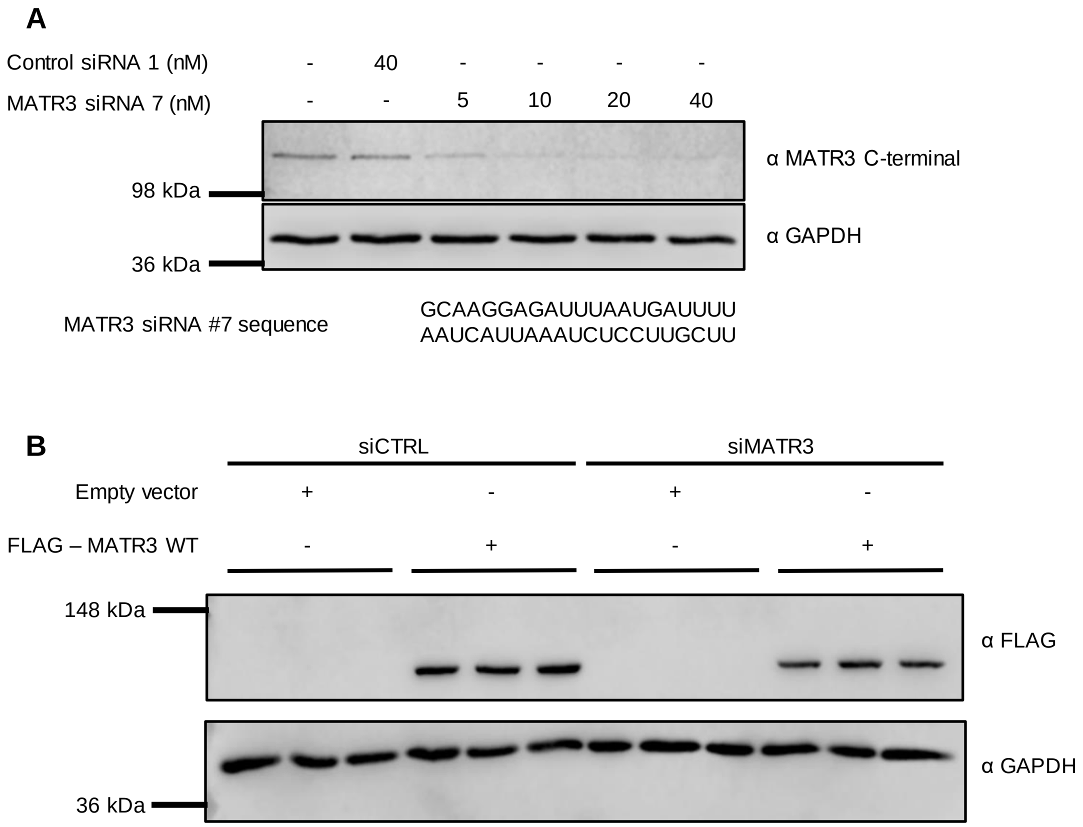


**Supplementary Figure 8. Optimization of MATR3 WT rescue experiment. A.** Dosage test for MATR3 siRNA #7. HeLa cells transfected with increasing amounts of MATR3 siRNA from 5 nM to 40 nM for 48 hours. The level of MATR3 knockdown was assessed by western blot analysis probing for MATR3 and comparing to GAPDH control. **B.** Western blot showing expression of MATR3 from the pcDNA–FLAG–MATR3 construct.

#### Figure S9

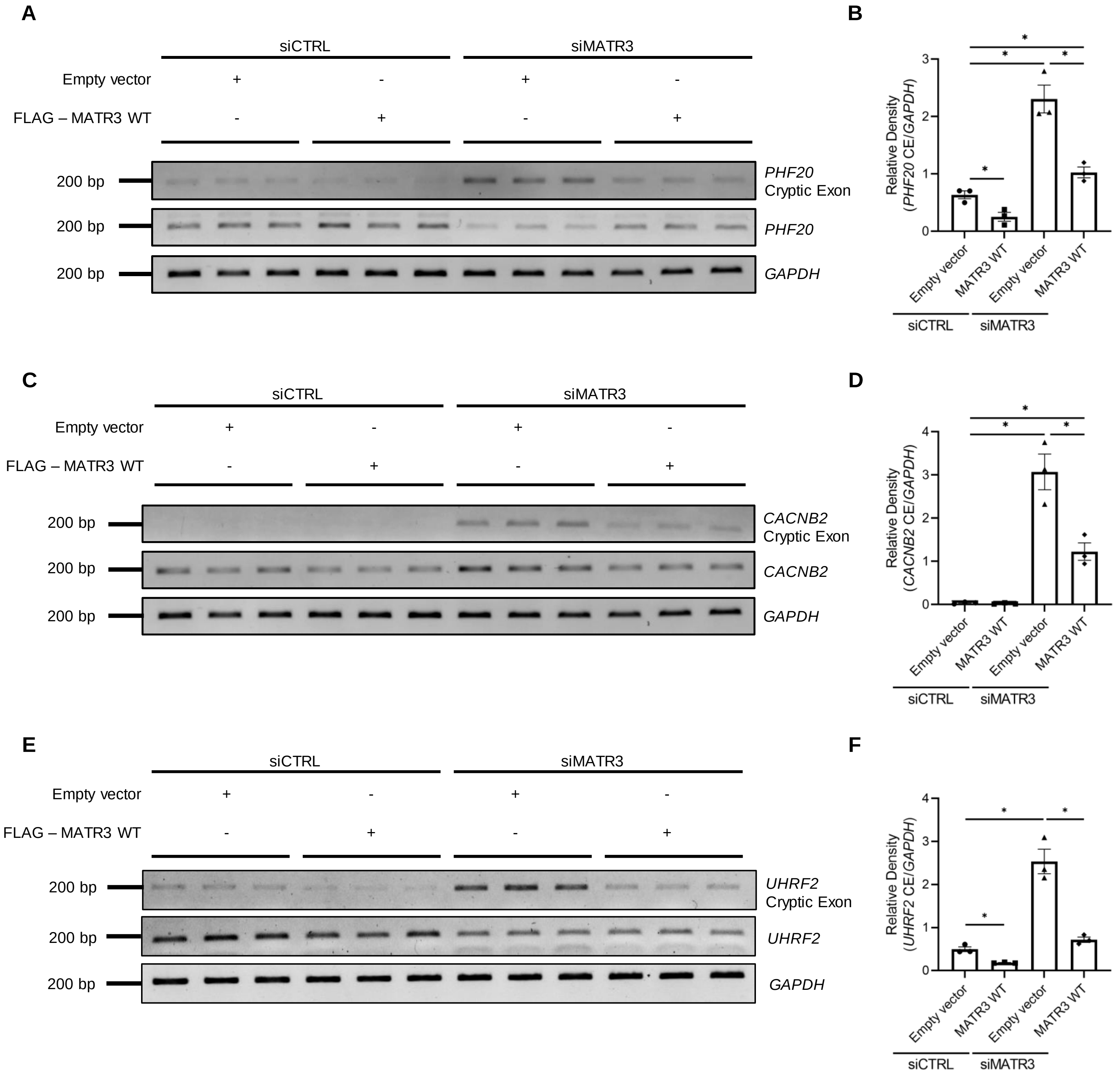


**Supplementary Figure 9. RT-PCR confirms cryptic exon inclusion in *PHF20, CACNB2,* and *UHRF2* upon loss of MATR3 was rescued by MATR3 WT. A.** Representative gels showing RT-PCR from total RNA extracted from HeLa cells transfected with siRNA and MATR3 vectors showing *GAPDH* and *PHF20* and **C.** *GAPDH* and *CACNB2* and **E.** *UHRF2* and *GAPDH.* Quantification of cryptic exon (CE) inclusion in **B.** *PHF20,* **D.** *CACNB2,* and **F.** *UHRF2* normalized to *GAPDH* (n=3, bar heights depict mean ± SEM, with each datapoint representing a biological replicate, significance determined by Welch's t-test, * $p \leq 0.05$).

#### Figure S10

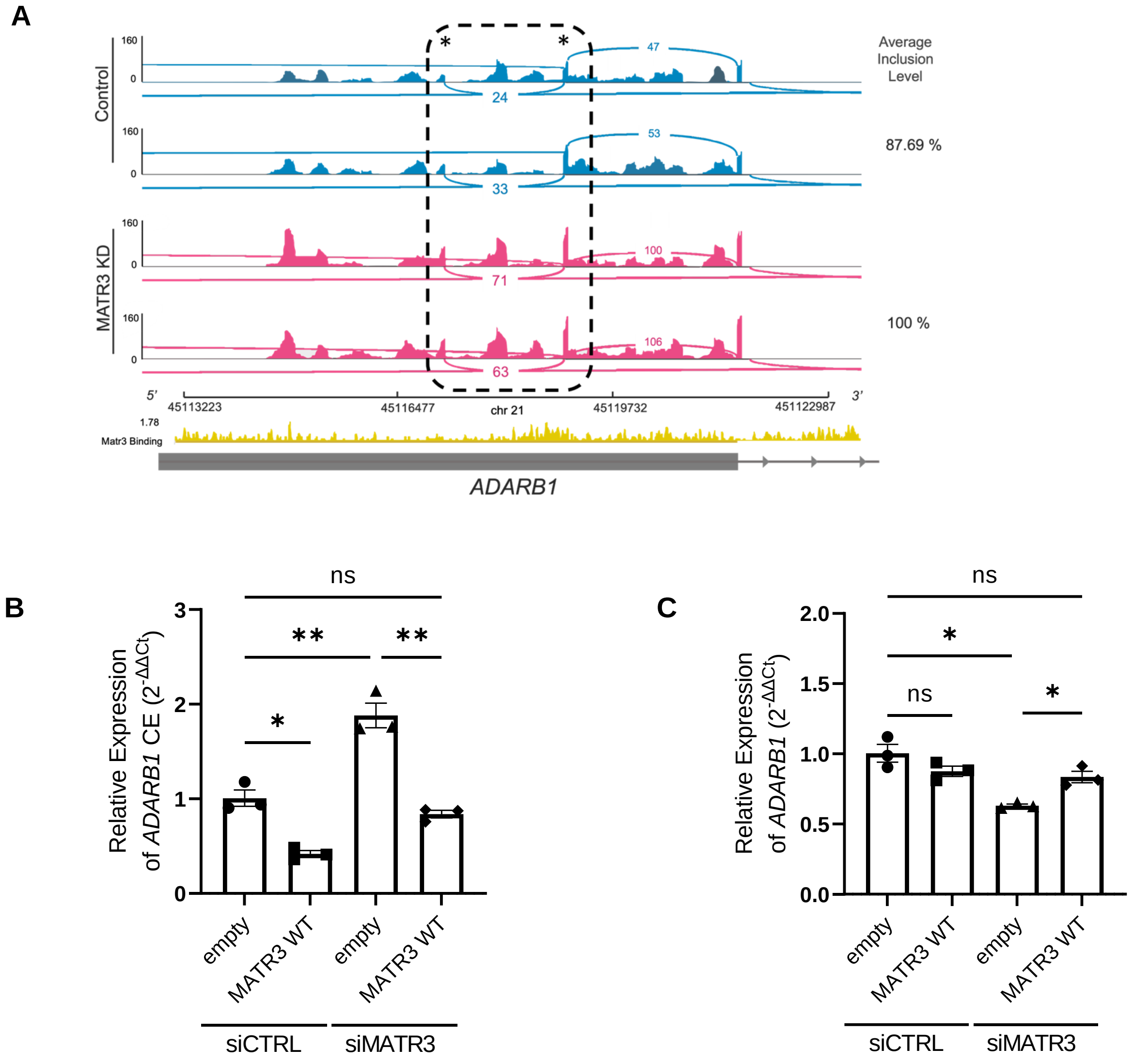


**Supplementary Figure 10. Cryptic splicing events upon MATR3 knockdown in *ADARB1*. A.** Sashimi plot generated using IGV genome browser illustrating the read coverage for two biological replicates of control (blue) and *MATR3* knockdown (pink) for a region of *ADARB1* transcribed from left to right. The bar graph represents read depth, and the arcs represent the number of reads connecting each splice junction, with junction coverage minimum set to 10. The region within black dashed lines represents the cryptic junction of interest with the cryptic exons marked by asterisks, and inclusion levels of the cryptic exon within this region are shown as percentages to the right of the tracks. MATR3 binding analyzed from CLIP-Seq data from Coelho *et al*., 2015 is shown in yellow. **B.** Relative gene expression (ΔΔCt analysis) of *ADARB1* mRNA containing the cryptic exon in HeLa cells doubly transfected with either nontargeting control siRNA (siCTRL) or siRNA targeting MATR3 (siMATR3) and FLAG-GFP or FLAG-MATR3 wildtype (WT) as measured by quantitative PCR (n=3, bar heights depict mean ± SEM, with each datapoint representing a biological replicate, significance determined by Welch's t-test, ns $p > 0.05$, ** $p \leq 0.01$, *** $p \leq 0.001$).

#### Figure S11

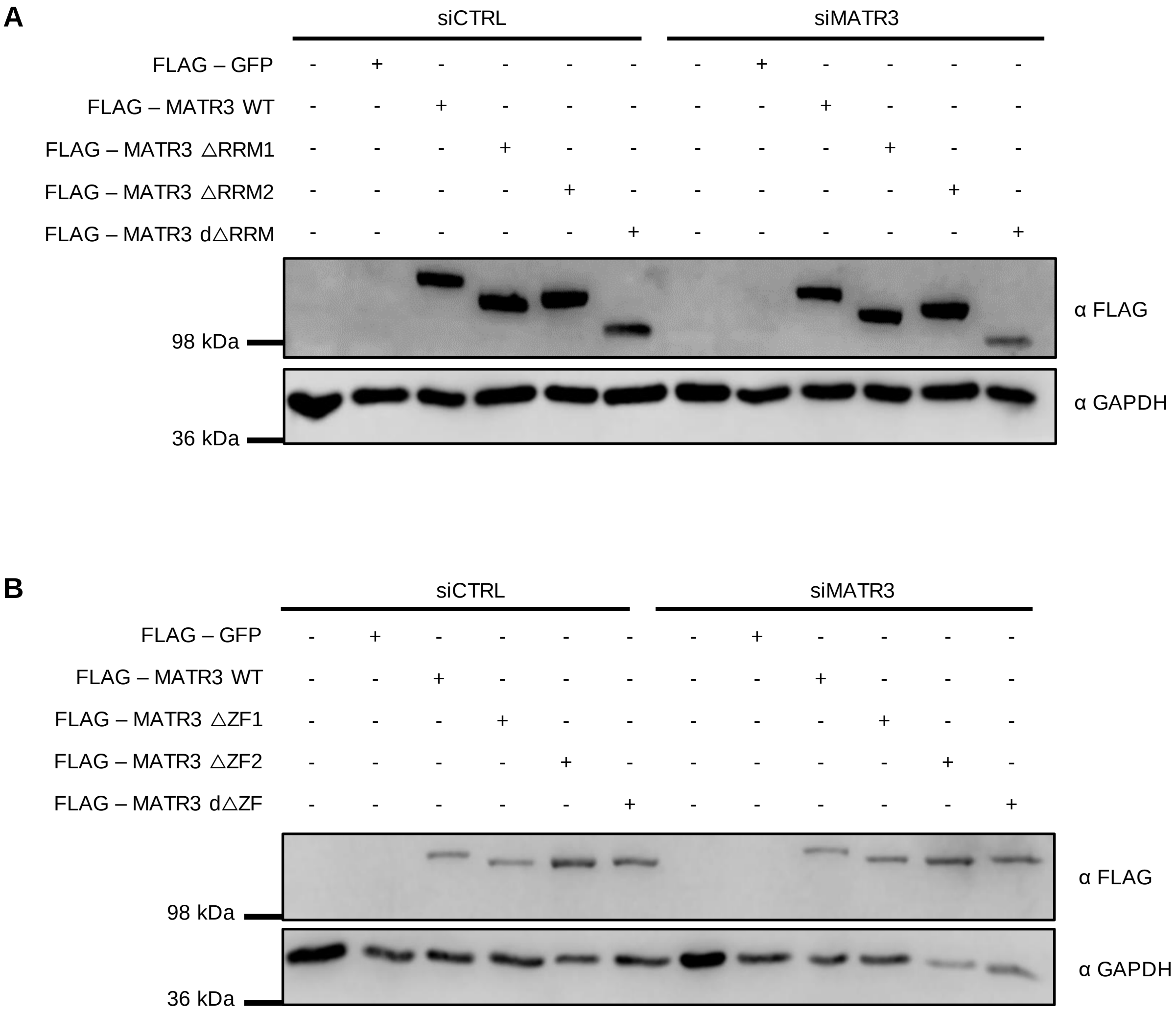


**Supplementary Figure 11. Expression of MATR3 domain deletion mutants. A.** Western blot showing expression of MATR3 RRM domain deletion mutants. **B.** Western blot showing expression of MATR3 ZF domain deletion mutants.

#### Figure S12

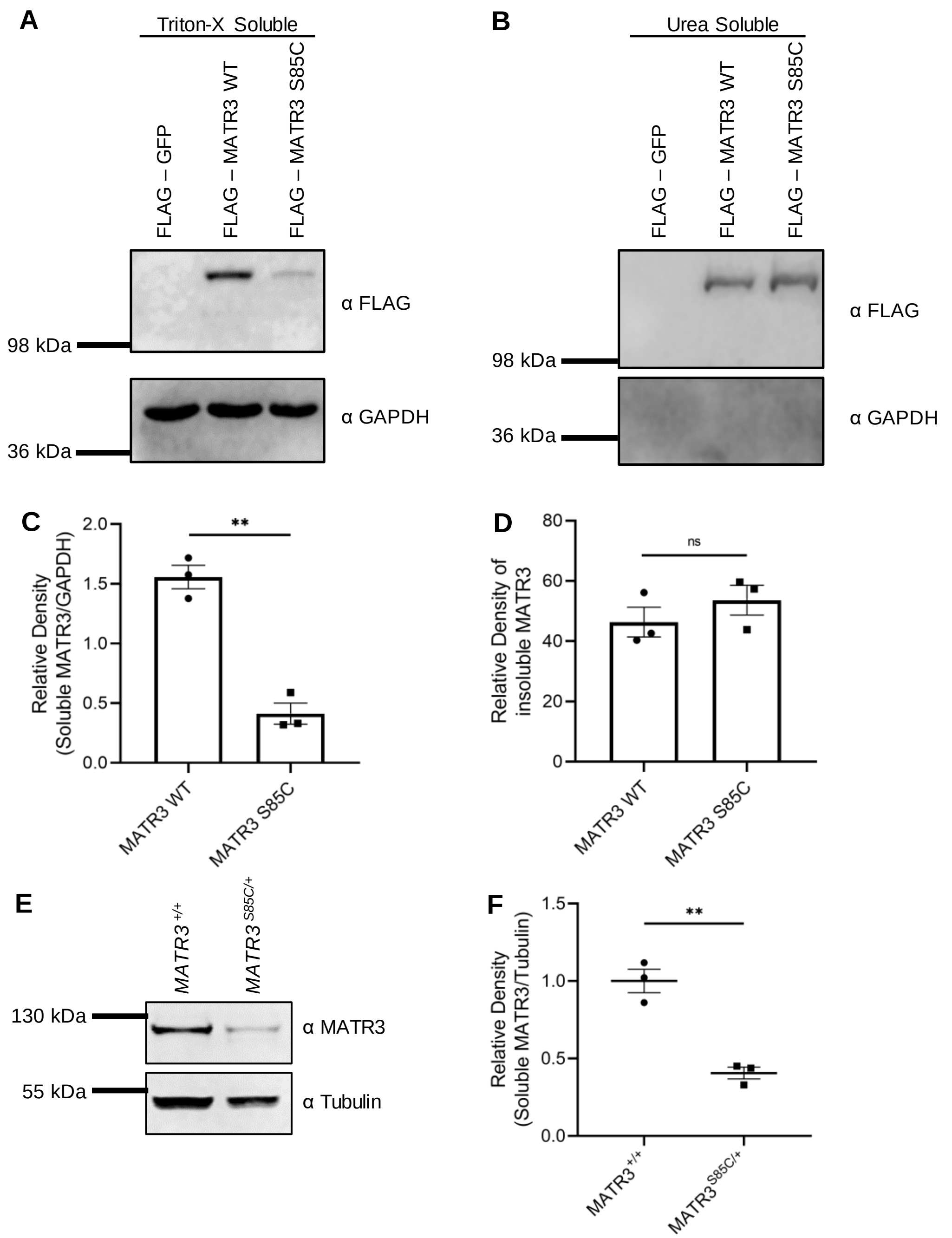


**Supplementary Figure 12. Soluble and insoluble fractionation of MATR3 S85C shows it is less soluble than MATR3 WT.**
**A.** Western blot showing Triton-X soluble (soluble fraction) protein levels of MATR3 WT and MATR3 S85C. **B.** Western blot showing urea soluble (insoluble fraction) protein levels of MATR3 WT and MATR3 S85C. **C.** Quantification of soluble MATR3 levels normalized to GAPDH (n=3, bar heights depict mean ± SEM, with each datapoint representing a biological replicate, a total of 3 independent experiments was performed, significance determined by Welch's t-test, ** $p \leq 0.01$). **D.** Quantification of insoluble MATR3 levels (n=3, bar heights depict mean ± SEM, with each datapoint representing a biological replicate, a total of 3 independent experiments was performed, significance determined by Welch's t-test, ** $p \leq 0.01$). **E.** Representative western blots showing soluble MATR3 and Tubulin protein levels in the differentiated *MATR3*$^{S85C/+}$ and isogenic control neurons. **F.** Quantitative graph depicting the decrease in soluble MATR3 protein in *MATR3*$^{S85C/+}$ neurons as compared to isogenic control (n=3, significance determined by Welch's t-test, ** $p \leq 0.001$).

**Figure S13**

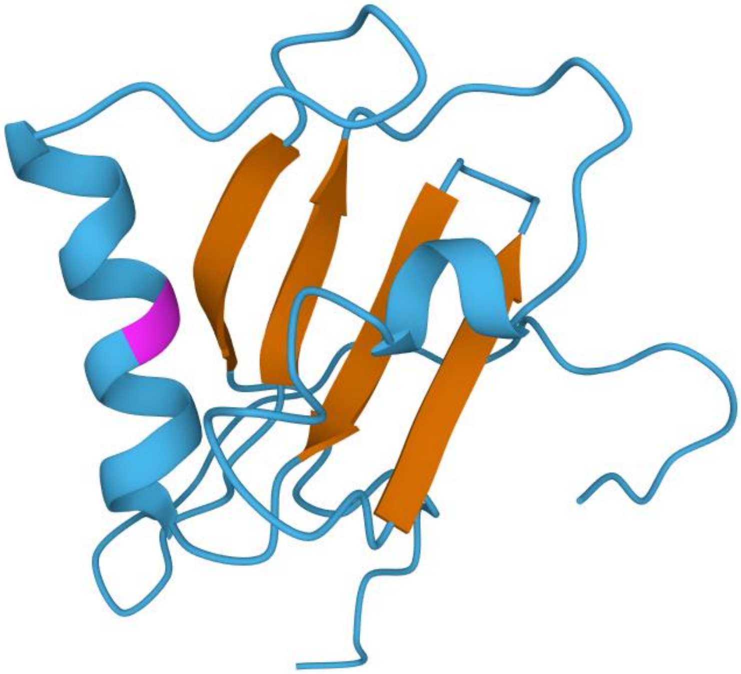

**Supplementary Figure 13. NMR structure of the RRM2 domain of MATR3.** The beta strands are shown in orange while the alpha helices and remainder of the domain are shown in blue. The location of the M548 residue is shown in magenta. Image from the RCSB PDB (rcsb.org) of PDB ID 1X4D (He, F., Kuwasako, K., Takizawa, M., Takahashi, M., Tsuda, K., Nagata, T., Watanabe, S., Tanaka, A., Kobayashi, N., Kigawa, T., et al. (2021). 1H, 13C and 15N resonance assignments and solution structures of the two RRM domains of Matrin-3. Biomolecular NMR assignments, 10.1007/s12104-021-10057-0)
